# Telomeres are shorter in wild *Saccharomyces cerevisiae* isolates than in domesticated ones

**DOI:** 10.1101/2022.02.03.478944

**Authors:** Melania D’Angiolo, Jia-Xing Yue, Matteo De Chiara, Benjamin P. Barré, Marie-Josèphe Giraud Panis, Eric Gilson, Gianni Liti

## Abstract

Telomeres are ribonucleoproteins that cap chromosome-ends and their DNA length is controlled by counteracting elongation and shortening processes. The budding yeast *Saccharomyces cerevisiae* has been a leading model to study telomere DNA length control and dynamics. Its telomeric DNA is maintained at a length that slightly varies between laboratory strains, but little is known about its variation at the species level. The recent publication of the genomes of over 1000 *S. cerevisiae* strains enabled us to explore telomere DNA length variation at an unprecedented scale. Here, we developed a bioinformatic pipeline (Y^ea^ISTY) to estimate telomere DNA length from whole-genome-sequences and applied it to the sequenced 1011 *S. cerevisiae* collection. Our results revealed broad natural telomere DNA length variation among the isolates. Notably, telomere DNA length is shorter in those derived from wild rather than domesticated environments. Wild isolates are enriched in loss-of-function mutations in genes known to regulate telomere DNA length and the return of domesticated yeasts to a wild habitat coincides with shorter telomeres. Moreover, telomere DNA length variation is associated with mitochondrial metabolism, and this association is driven by wild strains. Overall, these findings suggest that budding yeasts’ telomere DNA length regulation might be shaped by ecological life-styles.

## Introduction

Telomeres are ribonucleoprotein structures located at the ends of chromosomes (Blackburn & Gall, 1978). They comprise tandem repeats of a DNA sequence whose motif and length vary among species (Fulnečková et al., 2013; Monaghan, 2010). They maintain chromosome structure by distinguishing natural chromosome-ends from accidental double-stranded breaks (DSBs) and by contributing to chromosome segregation during cell division. Telomere DNA length (TL) is determined by the opposite actions of elongation and degradation pathways: elongation is ensured by telomerase, a reverse transcriptase that uses RNA as a template (Greider & Blackburn, 1985, 1987), or by alternative lengthening of telomeres (ALT) recombination pathways (Bryan et al., 1995; Lundblad & Blackburn, 1993; Teng & Zakian, 1999); degradation is linked to the semiconservative replication of DNA extremities coupled to specific nuclease activities (Lingner et al., 1995; Olovnikov, 1973; Watson, 1972). In mammalian somatic cells, telomerase is not or only weakly expressed and telomeres shorten at each cell division (Gilson & Géli, 2007). By contrast, telomerase is constitutively expressed in the budding yeast *Saccharomyces cerevisiae* allowing telomeric DNA to be maintained at an equilibrium length (Marcand et al., 1999).

TL differs broadly among species. Human telomeres can range between 5 and 15 kb while mice can have extremely long telomeres (even >100 kb) (Monaghan, 2010). In comparison, yeast telomeres are relatively short at around 300 bp (Teixeira & Gilson, 2005). Yeast telomeric sequences are often preceded by the so-called telomere-associated sequences (TAS). These comprise repetitive sequences like the X and Y’ elements, and interstitial telomeric sequences (ITS), which are located between them (Louis, 1995; Wellinger & Zakian, 2012). TL is a complex trait: in budding yeast it is modulated by more than 400 Telomere Length Maintenance (TLM) genes that were identified in large-scale systematic genetic screens (Askree et al., 2004; Gatbonton et al., 2006; Harari & Kupiec, 2014; Ungar et al., 2009), and by various environmental factors, including heat, caffeine and ethanol (Harari et al., 2020; Harari & Kupiec, 2018b; Kupiec, 2014; Romano et al., 2013). TL also varies among individuals of the same species, including yeasts, nematodes, plants, mammals and birds (Cook et al., 2016; Foley et al., 2020; Fulcher et al., 2014; Hansen et al., 2016; Heidinger et al., 2012; Liti et al., 2009; Zijlmans et al., 1997).

TL can be inferred from next-generation sequencing data by estimating the fraction of reads containing a sufficiently high number of telomeric DNA repeated sequences. This approach was implemented for human genomes containing regular telomeric DNA repeats (T_2_AG_3_)_n_ (Ding et al., 2014; Hakobyan et al., 2016; Lee et al., 2017; Nersisyan & Arakelyan, 2015) as well as for *Saccharomyces cerevisiae*, which contains irregular telomeric DNA repeated sequences (TG1-3)n (Puddu et al., 2019). These methods are now broadly applied (Barthel et al., 2017; Cook et al., 2016; Hakobyan et al., 2016; Nersisyan et al., 2019; Taub et al., 2020).

In humans, TL variation has been associated with multiple pathological conditions (Aviv & Shay, 2018) and to selection to attenuate the risk of cancer (Hansen et al., 2016; Mangino et al., 2015). In birds, TL has been used to predict key life-history traits like growth, reproduction and lifespan, and individuals with shorter TL have a lower life expectancy (Haussmann et al., 2005; Heidinger et al., 2012). Examples of extreme TL variation among different subpopulations have also been reported in yeast, including *S. cerevisiae* and its closest wild relative *S. paradoxus* (Liti, Haricharan, et al., 2009). This raises the question whether TL variation plays a role in the adaptation of a strain to its environment or if it derives from genetic drift. The lack of an adequate number of strains, especially the ones isolated from wild habitats, and experimental follow-up, has so far prevented this question from being addressed.

The recent publication of the genomes and phenomes of more than 1000 *S. cerevisiae* isolates (De Chiara, Barré, et al., 2020; De Chiara, Friedrich, et al., 2020; Peter et al., 2018) enabled us to explore TL variation at an unprecedented scale. Here, we report the first comprehensive analysis of TL variation in yeast strains isolated from a wide range of ecological niches and geographical areas, including domesticated and wild environments. For this purpose, we developed a bioinformatic pipeline that estimates the average TL, interstitial telomeric sequences content (ITS) and Y’ copy number from whole-genome sequencing data. We described TL variation across 26 well-defined lineages and identified new candidate genetic variants involved in TLM. We compared TL with a series of phenotypes including growth using different carbon and nitrogen sources, mitochondrial metabolism, sporulation capacity and chronological lifespan (CLS). The results revealed different TL regulation patterns in wild and domesticated yeasts that correlate with their mitochondrial functions.

## Results

### *S. cerevisiae* telomere analysis from whole genome sequences

We developed Y^ea^ISTY (<u>Y</u>east <u>ITS</u>, <u>T</u>elomeres and <u>Y’</u> elements estimator) to estimate TL, ITS content and Y’ copy number from yeast whole genome sequencing data (**Methods**). The performance of Y^ea^ISTY has been extensively benchmarked against simulated and experimental datasets and its running parameters have been optimized. The Y^ea^ISTY workflow begins with a screening of Illumina paired-end reads to retain the ones containing telomeric sequences (**Fig. 1a and Supplementary Fig. 1a**). Reads are retained if they contain a stretch of telomeric repeats (either C{1,3}A or TG{1,3}) longer or equal to 40 bp with only one gap of maximum 2 nucleotides allowed. Since telomeric and ITS-derived reads are virtually impossible to distinguish from each other based solely on their sequences, we introduced an additional step consisting in mapping all the reads to a modified reference genome in which all repetitive sequences, including telomeres, ITS and Y’ elements, have been masked. Moreover, a representative Y’ element was appended to this reference genome as an additional chromosome entry. The presence of a single representative Y’ element solves the problem of spurious mapping, provides a criterion to distinguish telomeric and ITS reads and enables us to reliably estimate Y’ elements copy number. In fact, the mapping position of the paired reads and their association with internal genomic regions or Y’ elements enable us to infer whether the read comes from a telomere or a ITS. ITS reads are recognized based on the following assumptions: i) if a read contains a stretch of C{1,3}A or TG{1,3} repeats and its pair maps on a non-subtelomeric part of the genome; ii) if a read contains a stretch of TG{1,3} repeats and its pair maps on the representative Y’ element. ITS reads are further classified as non-Y’-associated or Y’-associated if they fit the first or the second criterion, respectively. All the other retained reads from the first filtering step are considered as telomeric. For simplicity, we refer to Y’-associated ITS content as “ITS content”, while the amount of non-Y’-associated ITS is not further analysed in this study.

**Figure 1.**
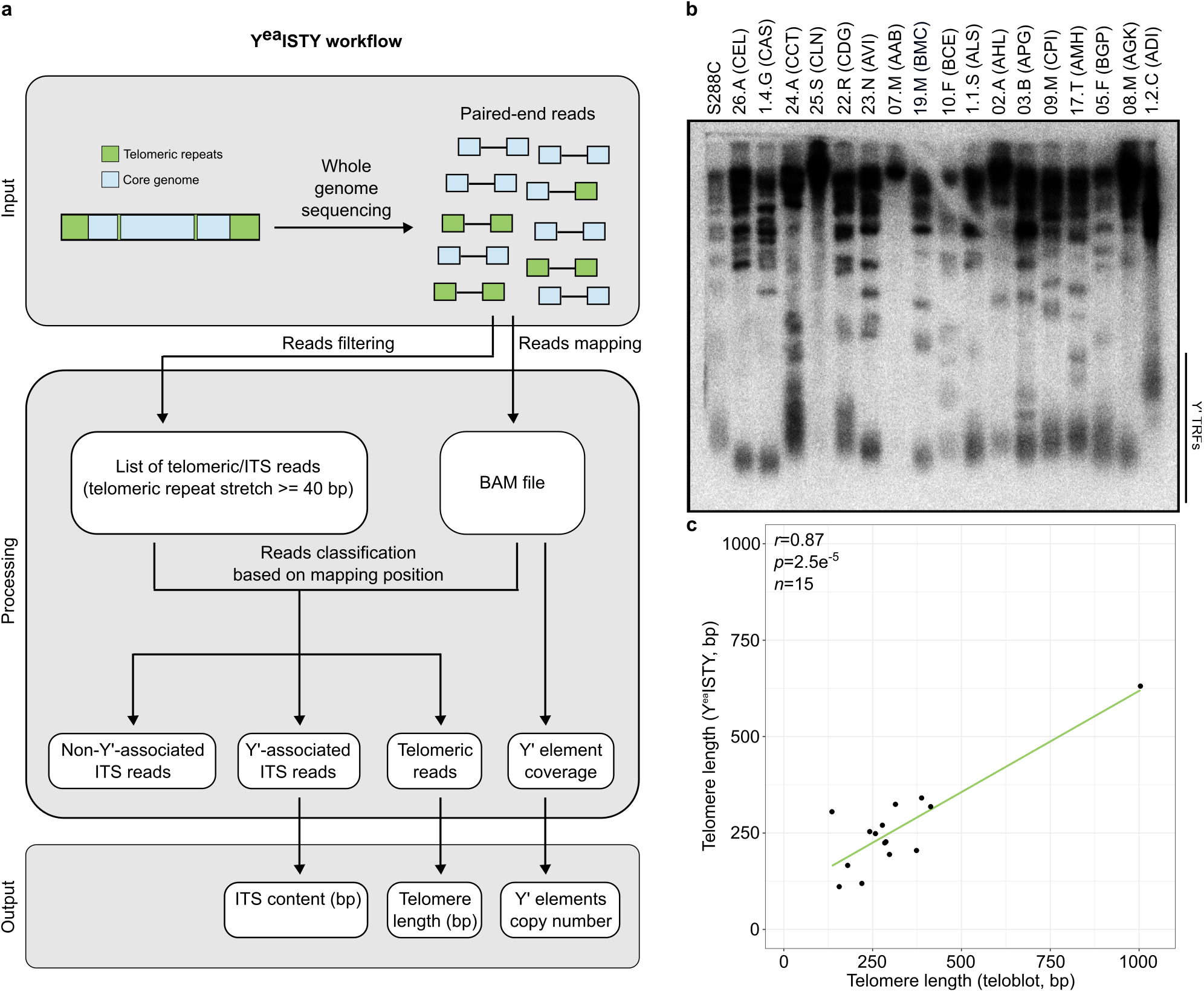
Overview of the Y^ea^ISTY workflow. **a**, Paired-end reads are scanned for telomeric repeats (either C{1,3}A or TG{1,3}). Reads containing a stretch of telomeric repeats longer or equal to 40 bp are retained. All the reads (retained and non-retained) are mapped to a modified reference genome. The mapping position of retained reads and their paired ones is used to classify them as ITS-derived or telomeric. The amount of ITS-derived and telomeric reads is then used to infer ITS content and telomere length, while coverage data are used to infer the copy number of Y’ elements. **b**, *Xho*I digestion of genomic DNA from representative strains of 17 *S. cerevisiae* lineages identified in Peter *et al.*, 2018. Genomic DNA is probed with radioactively-labelled telomeric TG_1–3_ repeats. The black line denotes terminal restriction fragments (TRFs) resulting from the digestion of a Y’ element. No TRFs were detected for the representatives of the Sake (25.S-CLN) and Mosaic beer (07.M-AAB) clades. **c**, Comparison of telomere length measured by teloblot and Y^ea^ISTY. Values in the plot are already corrected for the 4-fold Y^ea^ISTY underestimation bias. Despite we estimated TL of S288C from sequencing data derived from Yue *et al.*, 2017, we did not introduce this value in the plot as it derived from another sequencing batch and had a different underestimation rate. The Sake (CLN) and Mosaic beer (AAB) representatives were excluded from the comparison.

ITS content is calculated as *2*(_1_*∑^n^ s_ITS_*)/*c*, while TL is calculated as (*1∑^n^ s_TEL_*)/*32c*, where *n* is the number of reads assigned to each category (Y’-associated ITS or telomeric), *s* is the length of the telomeric repeats stretch contained in the reads assigned to each category, and *c* is the median coverage in regions with a GC content of 50-80%, which is similar to what is found in yeast telomeric repeats. ITS values are multiplied by 2 because we are only able to detect Y’-associated ITS reads containing the TG{1,3} motif, but we assume there will be the same amount of ITS reads containing the C{1,3}A motif. TL values are divided by 32 because this is the number of telomeres per haploid genome. In addition, Y’ elements copy number is estimated as *c^Y’^/c*, where *c^Y’^* is the median coverage of the representative Y’ element and *c* is the median coverage along the whole genome (**Fig. 1a, Supplementary Fig. 1a and Supplementary Discussion 1**).

We assessed the efficiency of Y^ea^ISTY by using the standard technique of Southern blotting (teloblot). We chose one representative strain for 17 clades of the *S. cerevisiae* collection and measured TL by teloblot. We correlated these results with the Y^ea^ISTY estimations and found an overall positive correlation (Pearson’s *r*=0.87, *p*=2.5e^-5^) (**Fig. 1b-c and Supplementary Table 1**). Two strains (25.S-CLN and 07.M-AAB) did not give any terminal restriction fragment (TRF) and were not included in the statistical analysis. One strain (1.2.C-ADI) had extremely long telomeres, in accordance with previous reports showing that this strain has atypical TL and massive amplification of ITS and Y’ elements (Bergstrom et al., 2014). However, TL resulting from Y^ea^ISTY was underestimated by 4-fold respect to the values of the teloblot. We therefore applied a 4-fold correction factor to the estimations of the *S. cerevisiae* collection. A detailed description of the implementation of Y^ea^ISTY’s algorithm is presented in supplementary discussion 1, while further benchmarking of Y^ea^ISTY on additional datasets is presented in supplementary discussions 2 and 3 (**Supplementary Fig. 1-4 and Supplementary Tables 2-4**). Taken together, these results show that Y^ea^ISTY estimations are reliable and can be used for further analyses.

### Global telomere length variation in the *S. cerevisiae* population

We applied Y^ea^ISTY to the 1011 *Saccharomyces cerevisiae* collection and estimated their TL. We retained 918 strains sequenced within the 1011 yeast genome project with comparable TL estimates and used these for further analyses (**Methods**). TL was normally distributed with a median of 236±96 bp, a value that is close to that of the S288C laboratory strain. Some isolates had either very short (minimum 41 bp) or very long (maximum 1210 bp) telomeres (**Supplementary Fig. 5a and Supplementary Table 5**). TL varied significantly among 26 previously described clades and three mosaic lineages (two-tailed Kruskal-Wallis test, *p*=9.154e^-7^) (Peter et al., 2018). TL was longer in the Alpechin and Mexican agave clades, where the median exceeded 300 bp, and it was shorter in the Malaysian and Ecuadorean clades, where it remained below 200 bp (**Fig. 2 and Supplementary Table 6**). The Wine European clade’s TL had a distribution comparable to that of the entire collection, ranging from 45 to 750 bp (median=238±101 bp), consistent with the largest clade sample size (~1/3 of the whole collection) and intra-clade substructure consisting of four subclades (Semi-wild, Clinical/Y’ amplification, Clinical/*S. boulardii*, Georgian) (**Supplementary Fig. 5b**). The median TL varied significantly among the Wine European subclades (two-tailed Kruskal-Wallis test, *p*=3.394e^-8^). It was highest in the Clinical/Y’ amplification subclade where it exceeded 600 bp, and lowest in the Clinical/*S. boulardii* subclade, where it remained below 200 bp (**Supplementary Fig. 5c and Supplementary Table 6**). Clinical/Y’ amplification strains are known to have aberrant TL and amplification of interstitial telomeric sequences (ITS) and Y’ elements (Bergstrom et al., 2014).

**Figure 2.**
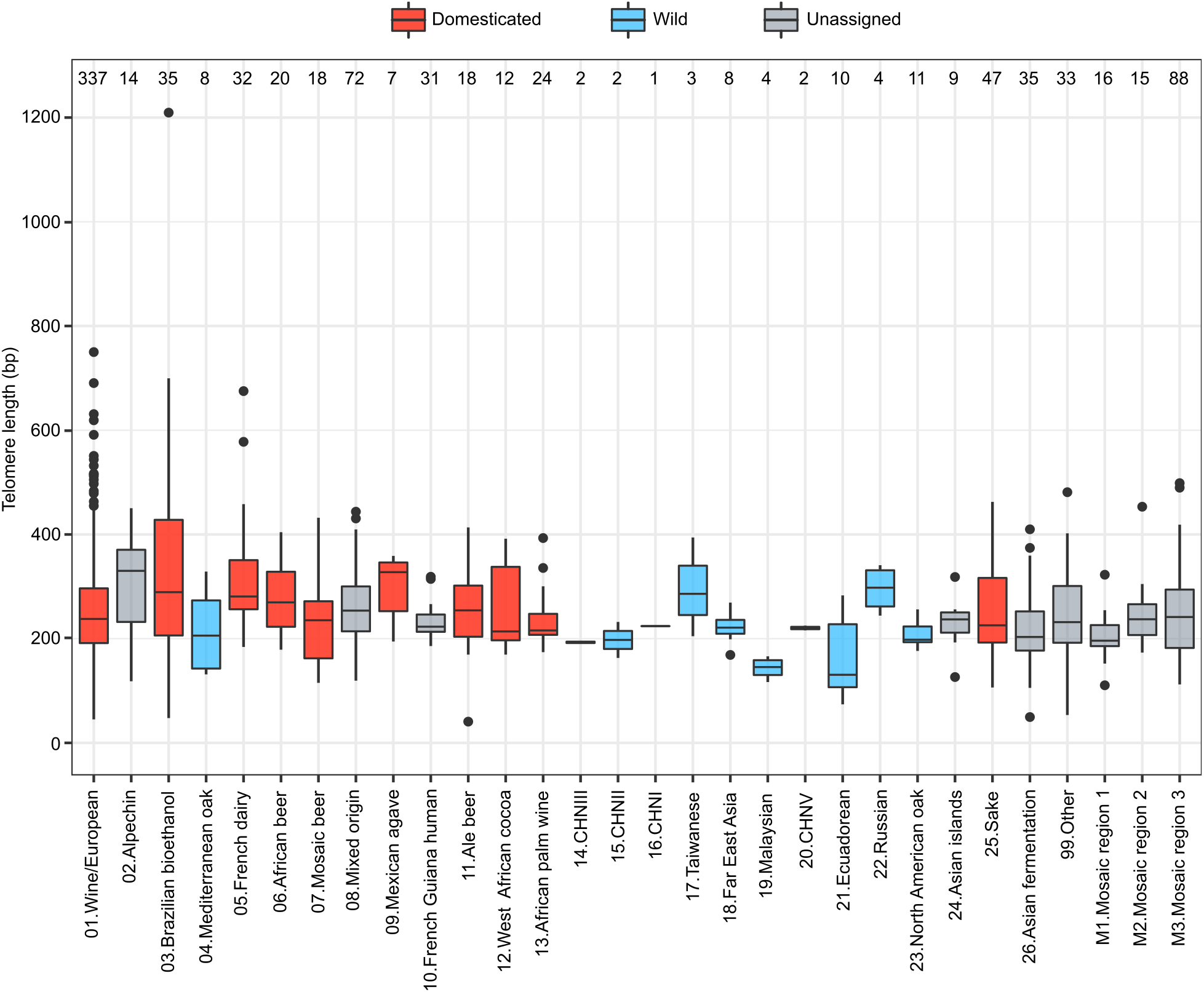
TL variation in the 918 *S. cerevisiae* collection. Telomere length of the phylogenetic lineages in the 918 *S. cerevisiae* collection (Peter *et al.*, 2018). Colours code represents the clade classification (domesticated, wild or unassigned), as reported in De Chiara, Barré *et al.*, 2020. Clades are assigned as wild or domesticated based on the environmental origin of the isolates dominating (>66%) in the clade. They were labelled “Unassigned” if the threshold criterion was not passed. The order of clades on the *x* axis is the same as in Peter *et al.*, 2018. In the box plots, horizontal lines denote the median, top and bottom hinges denote the IQR, whiskers denote maximum and minimum values within upper and lower hinges ± 1.5 × IQR. Numbers on top represent the number of isolates in each clade.

No variation in TL was detected among strains with variable ploidy and number of aneuploid chromosomes (two-tailed Kruskal-Wallis test, *p*=0.15 and *p*=0.76, respectively) (**Supplementary Fig. 5d-e**).

Finally, we measured the ITS content and Y’ copy number across the 918 isolates and found them to be highly variable but positively correlated with TL (Pearson’s *r*=0.43 and *p*<2.2e^-16^ between TL and ITS, *r*=0.39 and *p*<2.2e^-16^ between TL and Y’) and between themselves (Pearson’s *r*=0.81, *p*<2.2e^-16^) (**Supplementary Discussion 4, Supplementary Fig. 6-8 and Supplementary Tables 7-8**). The marked correlation between ITS and Y’ is consistent with the presence of short stretches of telomeric repeats between multiple Y’ copies (Louis & Haber, 1992). Of note, ITS and Y’ content in the Wine European subclades followed the same trend of TL, being more abundant in the Clinical/Y’ amplification subclade and less abundant in the Clinical/*S. boulardii* subclade. Overall, our analyses show that TL and ITS exhibit both inter- and intra-clade variation, but co-vary in the same direction.

### Domesticated and wild yeast isolates exhibit different TLs

Since domestication represents a crucial life-style shift in *Saccharomyces cerevisiae* (De Chiara, Barré, et al., 2020), we investigated whether it shaped TL. Isolates were previously classified as wild (*n*=55) or domesticated (*n*=550) based on the dominant origin (>66%) in their clade (De Chiara, Barré, et al., 2020; Peter et al., 2018). Domesticated isolates had significantly (two-tailed Wilcoxon test, *p*=9.296e^-5^) longer telomeres than wild ones (median=244±108 bp and 213±66 bp, respectively), with a median difference of ~30 bp (**Fig. 3a and Supplementary Table 5**). Consistently, TL was tendentially longer in domesticated clades than in wild ones (**Fig. 2**).

**Figure 3.**
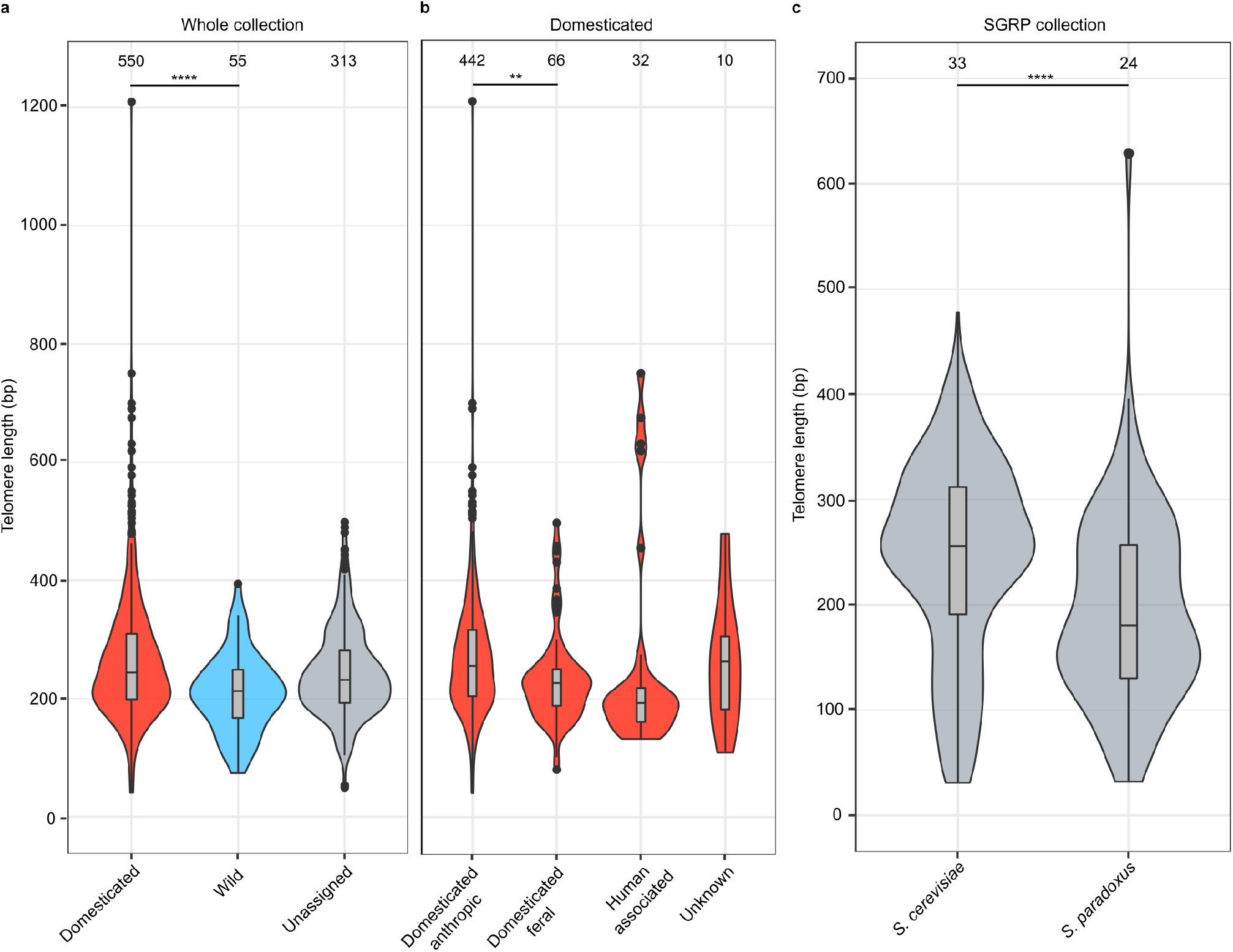
TL variation in domesticated and wild yeasts. **a**, Telomere length of domesticated, wild and unassigned isolates. Isolates classification is as in Fig. 2. **b**, Telomere length of domesticated isolates, divided in groups based on their substrate of isolation. “Domesticated-anthropic” are strains isolated from fermentation processes; “Domesticated-feral” are strains isolated from natural environments; “Human-associated” comprises strains isolated from the human body; “Unknown” comprises strains whose substrate of isolation is unknown. **c**, Telomere length of *S. cerevisiae* vs *S. paradoxus* strains from the *Saccharomyces* genome resequencing project (SGRP). Box plots within in the violin plots are as in Fig. 2. Numbers on top represent the number of isolates in each group. \**p* < 0.05, \*\**p* < 0.01, \*\*\**p* < 0.001, \*\*\*\**p* < 0.0001, ns=non-significant.

Within the domesticated group (*n*=550), most isolates were indeed derived from fermentation-related processes in line with the dominant origin of the clade. We refer to these strains as “domesticated-anthropic”, while others were isolated in nature and are defined here as “domesticated-feral” (**Supplementary Table 5**). Feral strains have gone through domestication but have then been re-dispersed into a natural environment. We compared TL between these two groups and found that feral isolates had a shorter median TL than anthropic ones (domesticated-anthropic=256±105 bp, domesticated-feral=227±80 bp, two-tailed Wilcoxon test, *p*=0.001) (**Fig. 3b**). This suggests that the long TL pattern is associated with ongoing domestication.

To further investigate the effects of domestication on TL, we compared the TL of *S. cerevisiae* to that of *S. paradoxus*, its closest related species which was never domesticated. We estimated the TL of *S. cerevisiae* (*n*=33) and *S. paradoxus* (*n*=24) strains from the *Saccharomyces* Genome Resequencing Project (SGRP) collection using already available Sanger sequencing data (Liti, Carter, et al., 2009). *S. cerevisiae* telomeres were longer than their *S. paradoxus* counterparts with a median difference of ~75 bp (*S. cerevisiae=256±93* bp, *S. paradoxus*=181±91 bp, two-tailed Wilcoxon test, *p*=3.96e^-12^). SGRP *S. cerevisiae* telomeres had lengths comparable to those of domesticated *S. cerevisiae* strains, while *S. paradoxus* telomeres had highly variable lengths with a median comparable to *S. cerevisiae* wild strains (**Fig. 3c, Supplementary Fig. 9 and Supplementary Table 7**). *S. paradoxus* TL is very different in its three continental subpopulations, with short telomeres in European isolates and very long ones in North American isolates despite all isolates being wild and isolated from similar ecological niches (Liti, Carter, et al., 2009; Liti, Haricharan, et al., 2009). Therefore, its overall resemblance to wild *S. cerevisiae* strains TL might derive by a sample bias toward European isolates and the high variance in the *S. paradoxus* group suggests that genetic drift in isolated subpopulations can also result in large TL variation.

We conclude that *S. cerevisiae* strains exhibit a broad distribution of TL patterns with medians that differ according to their life-styles: long for domesticated and short for wild strains. Remarkably, the domesticated pattern appears reversible, as exemplified in the feral isolates, showing that TL can change upon shifting life-style conditions and genetic drift.

### Natural genetic variants underlie TL variation

We searched for associations between common (minor allele frequency (MAF) > 5%) single nucleotide variants (SNV), copy number variants (CNV) and TL by conducting a genome-wide association study (GWAS) across 555 euploid diploid strains. We identified 20 variants (false discovery rate (FDR) corrected α=0.05), divided into 9 SNVs and 11 CNVs, including 2 CNVs encoded in the mitochondrial genome. Among the SNVs, 3 were in non-coding regions, while 6 were intragenic with 3 missense and 3 synonymous ones (**Fig. 4a and Supplementary Table 10**). Despite only one variant is in the intergenic region of a TLM gene previously identified in systematic genetic screens (*AHC2*) (Askree et al., 2004; Gatbonton et al., 2006; Ungar et al., 2009), 5 others are part of telomere length regulation pathways (*REV7, MIP6, PRI2, RPL7B, RNR3*). The remaining 14 variants underlie novel candidate TL regulators. The presence of intergenic hits suggests that part of TL regulation happens at the transcriptional level and these variants might act by regulating the expression of neighbouring TLM genes.

**Figure 4.**
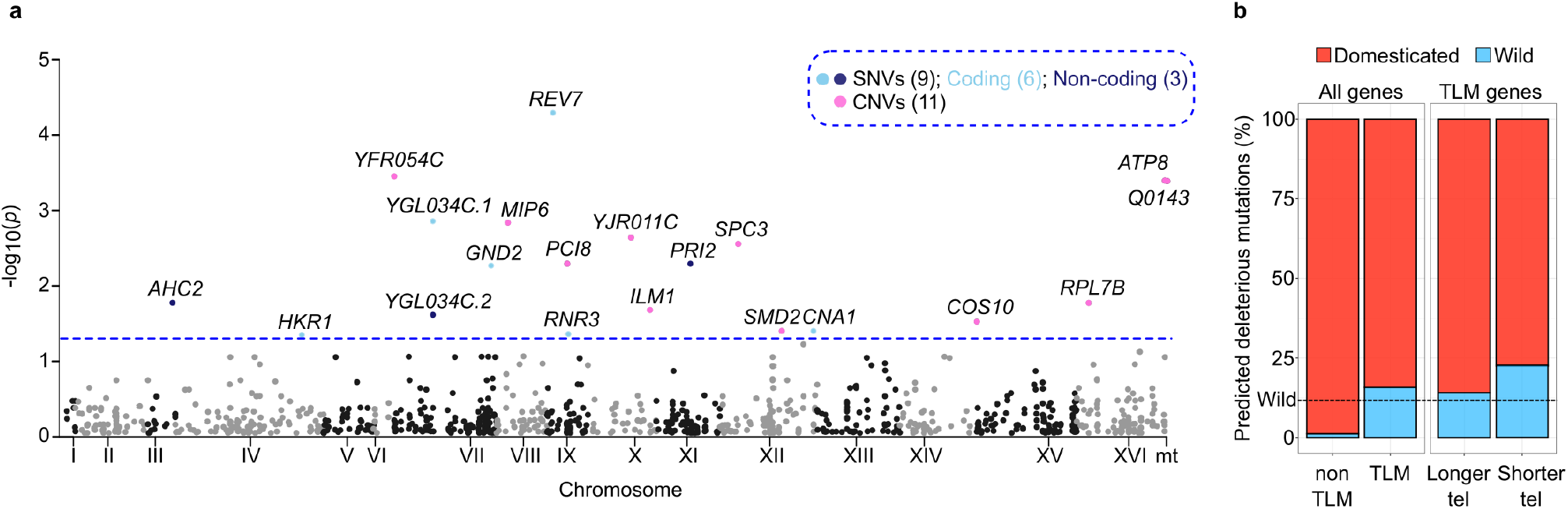
The genetic basis of TL variation. **a**, Manhattan plot showing the position of the GWAS variants across the genome. 20 variants are beyond the genome-wide significance threshold (*p*<0.05, blue dashed line). SNVs: single nucleotide variants; CNVs: copy number variants. Numbers in brackets denote the number of variants contained in that group. For simplicity, only the first 1000 tested variants are shown in the plot. **b**, Fraction of TLM and non-TLM genes carrying predicted loss-of-function mutations in euploid diploid domesticated (*n*=350) and wild (*n*=49) isolates (total *n*=399). Isolates classified as unassigned (*n*=156) were not considered in this analysis. TLM genes are further subdivided into the ones conferring shorter vs longer telomeres when deleted. The dashed line indicates the fraction of wild strains in the 399 *S. cerevisiae* collection (12%).

In most nuclear CNVs, gene copy number amplification was associated with shorter TL, except for *YFR054C*, and in most cases TL variation was stratified in the Clinical/*S. boulardii* subclade and the Ecuadorean clade, which carry multiple gene copies (**Supplementary Fig. 10a**). Similarly, both mitochondrial CNVs inversely correlated with TL. Since accurate CN estimation of individual genes in the mitochondrial genome is difficult, we checked the relationship between TL and mtDNA copy number estimated using a set of 3 genes with non-problematic mapping (De Chiara, Friedrich, et al., 2020). Although the association was not statistically significant, there was anticorrelation between the two variables (**Supplementary Table 11**).

In most of the SNVs (except *CNA1*), the allele with the lowest frequency in the population (minor) was associated with longer telomeres, but no clades emerged as the main drivers of TL variation. The *CNA1* minor allele was significantly enriched in domesticated clades (two-tailed *X^2^* test, *p*=7.68e^-5^) and associated with longer telomeres, therefore contributing to the TL difference between domesticated and wild isolates (**Supplementary Fig. 10b and Supplementary Table 10**). Overall, we estimated TL narrow-sense heritability to be 34%, with the significant GWAS variants explaining 11% of the total phenotypic variance of the population.

Rare genetic variants (MAF<5%) evade GWAS detection, so we investigated their effect by selecting likely loss of function (LOF) variants in TLM genes (*n*=383). TLM LOFs were significantly enriched in wild strains (two-tailed *X^2^* test, *p*=6.7e^-12^). Noteworthy, this enrichment was more pronounced in genes whose deletion causes telomere shortening (*p*=7.7e^-19^) rather than lengthening (*p*=0.001), in accordance with the trend of wild isolates to have shorter telomeres. This effect is specific to telomere length regulation, as LOFs in non-TLM genes (*n*=5648) showed the opposite trend, being depleted in wild strains and enriched in domesticated ones (**Fig. 4b and Supplementary Table 11**). Some of the TLM LOFs were private to one or few clades and could contribute to explain the extreme TL observed in those lineages (**Supplementary Discussion 5 and Supplementary Table 11**).

In conclusion, we detected multiple genetic variants and LOF mutations that contribute to the TL variation observed in the *S. cerevisiae* collection segregating within specific clades.

### Mitochondrial metabolism is associated with TL variation

We measured mitochondrial volume and activity in the euploid diploid strains (*n*=555) in either fermentative (YPD) or respiratory (YPEG) medium (**Supplementary Table 12**). Both mitochondrial activity and volume in YPD and YPEG were positively correlated (Pearson’s *r*=0.48, p<2.2e^-16^ for activity, *r*=0.62, p<2.2e^-16^ for volume, **Supplementary Fig. 11a**). While activity was higher in the glycerol-containing medium YPEG, in which fermentative metabolism is not active and respiration drives growth, volume was higher in YPD (two-tailed Wilcoxon test, *p*<2.2e^-16^ for activity, *p*<2.2e^-16^ for volume, **Supplementary Fig. 11b-c**).

Mitochondrial activity and volume varied significantly among the clades (two-tailed Kruskal-Wallis test, *p*<2.2e^-16^ for all the mitochondrial phenotypes; only mitochondrial activity in YPD is shown in fig. 5a for simplicity). Mitochondrial activity in YPD was particularly high in the Alpechin clade (**Fig. 5a**). Wild isolates had lower mitochondrial activity and volume in both YPD and YPEG as compared to domesticated ones (two-tailed Wilcoxon test, *p*=5.58e^-7^ and *p*=2.93e^-9^ for mitochondrial volume and activity in YPD, respectively; *p*=1.06e^-14^ and *p*<2.2e^-16^ for mitochondrial volume and activity in YPEG, respectively; **Supplementary Fig. 11d-e**). We then examined the mitochondrial phenotypes of the feral strains and found they had a lower mitochondrial volume than anthropic ones (two-tailed Wilcoxon test, *p*=0.0008 in YPD and *p*=0.0008 in YPEG). Mitochondrial activity was also lower in feral strains but not significantly (two-tailed Wilcoxon test, *p*=0.22 in YPD and *p*=0.30 in YPEG).

**Figure 5.**
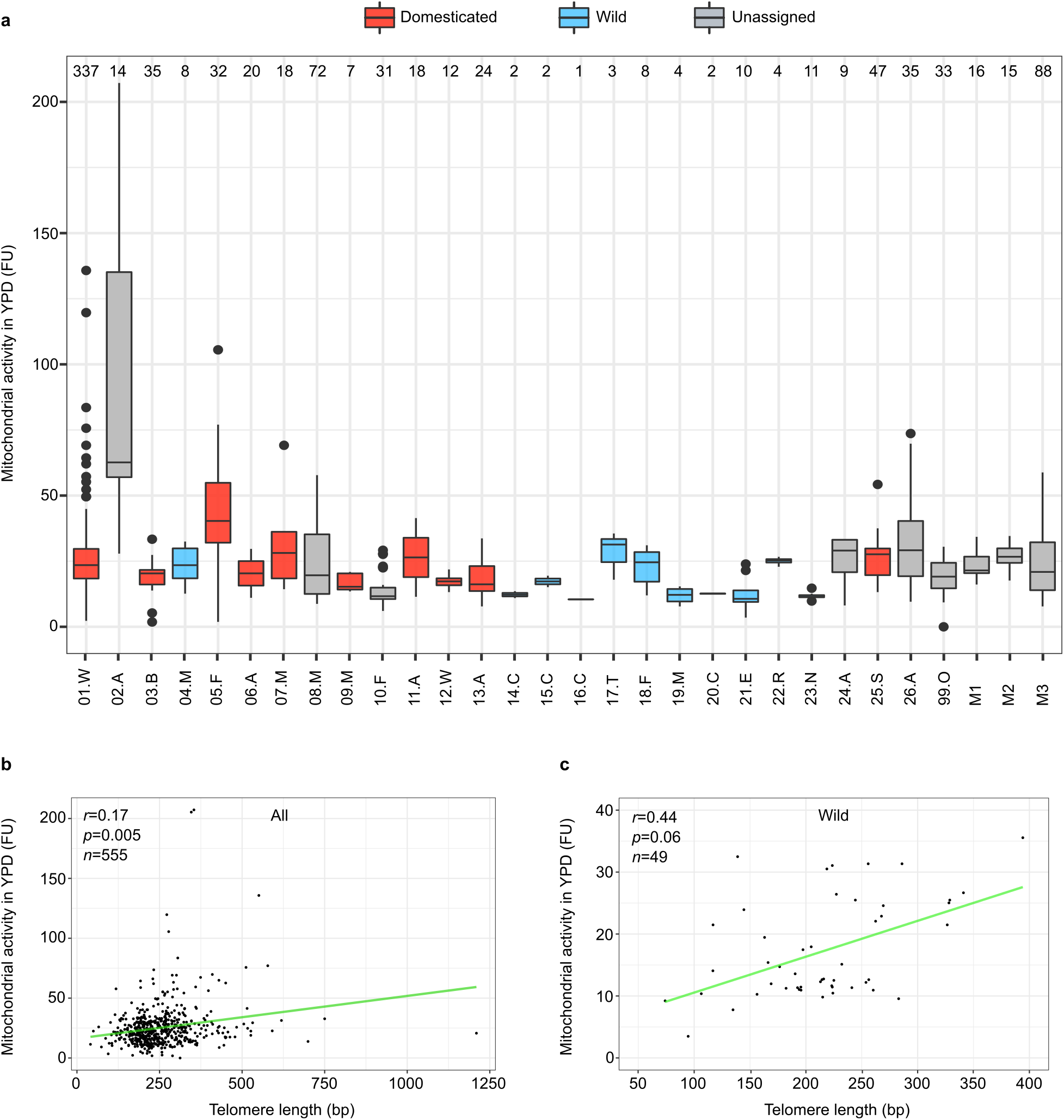
TL and mitochondrial phenotypes. **a**, Mitochondrial activity in YPD of the phylogenetic lineages in the 555 *S. cerevisiae* collection. Isolates classification is as in Fig. 2. Box plots are as in Fig. 2. FU=fluorescence units. **b-c**, Associations between mitochondrial activity in YPD and telomere length in the 555 *S. cerevisiae* collection (left panel) and in the subset of wild isolates (*n*=49) (right panel). Each point represents a single isolate and the line represents a linear regression function. The association is significant when considering the whole dataset (*n*=555), while it is borderline when considering only wild isolates (*n*=49). The other 3 mitochondrial phenotypes (mitochondrial activity and volume in YPD and YPEG) show similar trend but are not shown for simplicity, while the copy number of mitochondrial DNA negatively correlates with telomere length.

There were positive correlations of TL with mitochondrial volume and activity in both YPD and YPEG (Spearman’s *r*=0.21 and *p*=0.0002 for mitochondrial volume in YPEG, *r*=0.2 and *p*=0.0004 for mitochondrial activity in YPEG, *r*=0.14 and *p*=0.05 for mitochondrial volume in YPD, *r*=0.17 and *p*=0.005 for mitochondrial activity in YPD). Since domesticated and wild yeast isolates have different TL patterns, we checked their associations with the mitochondrial phenotypes both jointly and separately. The correlation coefficients of all mitochondrial phenotypes were higher when considering the wild isolates alone (Spearman’s *r*=0.31 for mitochondrial volume in YPEG, *r*=0.31 for mitochondrial activity in YPEG, *r*=0.31 for mitochondrial volume in YPD, *r*=0.44 for mitochondrial activity in YPD), although *p*-values failed to reach statistical significance in this setting because of the reduced sample size (**Fig. 5b-c**, only mitochondrial activity in YPD is shown for simplicity, **Supplementary Fig. 12 and Supplementary Table 12**). When switching to Pearson’s correlation, the coefficients were higher, indicating that TL and mitochondrial metabolism co-vary according to a linear model (**Supplementary Table 12**). In domesticated strains we observed the opposite trend, with correlation coefficients close to 0, consistent with wild strains being the main drivers of the association between TL and mitochondria.

Finally, we investigated the relationship between TL variation and a set of previously published phenotypes (*n*=151) including growth in different carbon and nitrogen sources, in the presence of chemical compounds and environmental challenges, reproductive capacity and chronological lifespan under standard (SDC) and caloric-restricted conditions (De Chiara, Barré, et al., 2020; De Chiara, Friedrich, et al., 2020; Peter et al., 2018). No significant correlations were found between TL and other phenotypes except for growth yield using raffinose as carbon source (2% and 8%) when considering all the isolates, resilience in the presence of salt (NaCl 1M) and growth yield using methionine as the only nitrogen source when considering only domesticated isolates, and chronological lifespan in standard (SDC) conditions (at days 7 and 21) when considering only wild isolates (**Supplementary Fig. 12 and Supplementary Table 13**).

Overall, these findings indicate that mitochondrial phenotypes mildly correlate with TL and this association is largely driven by wild isolates.

## Discussion

By estimating the telomeric DNA length (TL) and ITS content of 918 *Saccharomyces cerevisiae* isolates collected worldwide and associated with an extensive genomic and phenotyping dataset (De Chiara, Barré, et al., 2020; De Chiara, Friedrich, et al., 2020; Peter et al., 2018) we unveiled the species span of telomere length variation. We observed a wide spectrum of normally distributed TLs whose median value is close to what previously measured in the classical S288C laboratory strain. This variation is not explained by ploidy and aneuploidy and instead is associated with several punctuated genetic variants.

We analysed the association between TL and the ecological origin of the yeast isolates. Remarkably, we found that TL was shorter in wild strains than in domesticated ones, although the distributions within these two groups are broad. Noteworthy, within the domesticated group, the feral strains have shorter TL than anthropic ones, suggesting that the long TL pattern is associated with ongoing domestication while the return to wild-life is accompanied by TL shortening. The difference in TL between domesticated and wild strains further adds to the profound disparities in life-cycle and metabolism previously described, with wild strains growing better in stressful environments and uniformly having a more efficient sexual reproduction (De Chiara, Barré, et al., 2020). Our findings, together with a recent report showing that wild mammals have shorter TL than domesticated ones (Pepke & Eisenberg, 2021), unveil that domestication versus wild-life are important life-style traits that are associated with TL regulation.

These results raise the question of whether variations in TL underlie an adaptive strategy to different environments as it was previously proposed in various organisms (Casagrande & Hau, 2019; Jacome Burbano & Gilson, 2021; Young, 2018). On one hand, a wild habitat imposes yeasts to cope with fluctuating environments and short telomeres could help survival through the transcriptional regulation of subtelomeric regions, which are enriched in stress response, nutrient utilization and cell wall composition genes (Ai et al., 2002; Robyr et al., 2002; Smith et al., 2011; Stone & Pillus, 1996; Wyrick et al., 1999; Ye et al., 2014). In addition, short TL can lead to the release of transcription factors from telomeres, allowing them to bind and regulate their targets genome-wide (Buck & Lieb, 2006; Maillet et al., 1996; Martin et al., 1999; Platt et al., 2013). This hypothesis agrees with the shorter telomere DNA length observed in feral strains, suggesting that the return to wild settings coincides with short telomeres. On the other hand, long telomeres could be favoured in domesticated environments, where fermentative processes submit yeasts to previously unencountered stresses. Alternatively, TL may not be directly under selection but its variable pattern in domesticated/wild yeasts could be a corollary of the so-called “domestication syndrome” (De Chiara, Barré, et al., 2020). A neutral scenario dictated by genetic drift could also explain the broad TL variability observed within the two classes of strains. For example, the highly diverged Asian lineages have different TL despite sharing the same life-style, habitat and some life cycle traits, suggesting that TL was free to evolve after the split from their last common ancestor. The 1011 *S. cerevisiae* collection is underrepresented for wild isolates and a more extensive sampling of strains from natural environments will better elucidate the role of drift and/or selection in shaping TL. Further studies where fitness is measured in isogenic background with controlled TL and in culture conditions close to the wild and domestic environments are also needed to distinguish between these evolutionary scenarios.

Among a wide range of phenotypes tested, TL positively correlated with mitochondrial volume and activity while negatively correlated with mitochondrial DNA copy number. These results reinforce the view that telomeres and mitochondria are functionally connected (Nautiyal et al., 2002; Passos, Saretzki, & Von Zglinicki, 2007; Passos, Saretzki, Ahmed, et al., 2007; Qian et al., 2019; Robin et al., 2020; Sahin et al., 2011). Instead, TL was generally not associated with growth in various conditions, in line with previous results showing that artificially shortened/elongated telomeres do not affect yeast fitness in laboratory conditions (Harari et al., 2017; Harari & Kupiec, 2018a). Whether the association between mitochondria and telomeres in wild strains represents a fitness advantage is a possibility that warrants further investigation.

Overall, our analyses revealed extensive TL variation across a large population of *S. cerevisiae* isolates collected around the world, which can be explained, at least in part, by genetic variations, ecological origins and associations between telomere length and mitochondrial functions. The low narrow-sense heritability of TL in our population (34%) suggests that a substantial fraction of TL variation is affected by non-additive genetic factors. Furthermore, isogenic *S. cerevisiae* backgrounds grown under defined environmental conditions (e.g. ethanol, acetic acid and caffeine) adjust TL through the modulation of TLM protein levels to a magnitude that overlaps the variation detected in the 1011 collection, showing that this phenotype is highly plastic (Harari et al., 2013; Kupiec & Weisman, 2012; Romano et al., 2013; Ungar et al., 2011). Future studies will quantify the role of genes, environment and their interactions in controlling telomere length in natural populations.

## Methods

### A bioinformatic pipeline for estimating TL, ITS content and Y’ copy number

We developed Y^ea^ISTY (<u>Y</u>east <u>ITS</u>, <u>T</u>elomeres and <u>Y</u>’ elements estimator) in order to estimate TL, ITS content and Y’ copy number in the 1011 *S. cerevisiae* collection.

We set a threshold of 40 bp for the minimum length of the telomeric repeats stretch contained in the reads, and ran Y^ea^ISTY with the following command line:

~~~
perl find_motif_in_reads.pl -i $SAMPLE.fasta -m motif.txt -o $SAMPLE.fasta.readscan -l $SAMPLE.fasta.readlist -c INT
~~~

where “i” is the input file (in FASTA format), “m” is a file containing the telomeric motifs for pattern matching (C{1,3}A, TG{1,3}), “o” is an output file containing a list of the reads classified as telomeric or ITS-derived and the position and length of their telomeric motifs, “l” is another output file containing the names of the reads classified as telomeric or ITS-derived, and “c” represents the minimum number of bp covered by telomeric motifs that must be contained in a read in order to classify it as telomeric or ITS-derived. This value can be set by the user and must be an integer number (INT). In this study, we set “c” as 40. Of note, the file containing the motifs can be modified to search for any short repeat, meaning that Y^ea^ISTY can be used to estimate the content of any type of short repetitive sequence in a sample.

Subsequently, we performed mapping on a modified SGD reference genome in which we masked all repetitive sequences, including telomeres, ITS and Y’ elements, by BWA (version 0.7.12) (Li & Durbin, 2009). The list of repetitive sequences in the SGD genome was generated and masked using RepeatMasker (http://www.repeatmasker.org/). A representative, long version Y’ element was appended to this reference genome as an additional chromosome entry. The representative Y’ element was chosen to be the one from TEL09L, as previously described (Yue et al., 2017).

After applying the pipeline, 93 isolates stood out from the rest because they exhibited extremely high TL. These 93 isolates were sequenced in previous studies. Given that the results were not comparable to the rest of the collection, we excluded these isolates from this study and only considered the remaining 918 strains.

We observed an underestimation rate of 4- and 8-fold for TL and ITS content, respectively, as compared to the average values measured by teloblot and the annotation of the 12 genome assemblies. We adjusted this bias by increasing TL and ITS content of 4- and 8-fold, respectively, in the estimations of the 1011 *S. cerevisiae* collection. Values reported in supplementary table 5 are already adjusted. Estimates from the other datasets used in this study varied around a different average value respect to the one of the 1011 *S. cerevisiae* collection, indicating that the underestimation rate differs across datasets and therefore estimates are not directly comparable between datasets, but only within them (Puddu et al., 2019; Tattini et al., 2019). Since we did not perform teloblots of representative strains from all the other datasets, their exact underestimation rates remained unknown and their estimates were not adjusted. The exact reasons of this variation are unknown and might depend on the combination of the sequencing technology, library preparation methods and coverage. While relative values can still indicate whether one strain carries longer/shorter telomeres than another one, a comparison with already known TL and ITS is needed to determine the exact underestimation rate of each dataset and obtain absolute TL and ITS estimates.

Y^ea^ISTY constitutes a valid tool to estimate TL, ITS content and Y’ copy number from whole genome sequencing data and it is freely available through github (https://github.com/mdangiolo89/Different-telomere-length-patterns-between-domestic-and-wild-S.-cerevisiae-isolates).

### Simulations of sequencing runs from genomes carrying synthetic telomeres

To estimate the efficiency of Y^ea^ISTY, we applied it to artificially constructed datasets, consisting of simulated sequencing runs from genomes carrying synthetic telomeres of known length. We masked the native telomeres of the 12 previously described genome assemblies by using the program maskfasta contained in the suite bedtools (version 2.17.0) (Quinlan & Hall, 2010). Subsequently, we replaced the masked native telomeres with 14 synthetic telomeres of known length, consisting of yeast telomeric TG{1,3} or C{1,3}A repeats. We created 168 (12 x 14) synthetic genomes in each of which single telomeres were all identical to each other. The lengths of the synthetic telomeres had the following values: 18, 49, 77, 99, 135, 174, 208, 296, 360, 416, 504, 553, 602, 651 bp. ITS were not modified. Illumina reads were then simulated from the 168 synthetic genomes using the software dwgsim (https://github.com/nh13/DWGSIM) with the following parameters: type of read=paired-end, read length=100 bp, sequencing error rate=0%, insert size=300 bp, coverage=30.

### ITS content, Y’ copy number and TL estimation in the yeast population reference panel

We estimated TL, ITS content and Y’ copy number in 7 *S. cerevisiae* and 5 *S. paradoxus* strains sequenced with Pacbio technology and whose complete genome assemblies were already available as part of the yeast population reference panel (YPRP) (Yue et al., 2017). We estimated the frequency and distribution of stretches of telomeric repeats in the 12 genome assemblies by using custom Perl scripts. We merged the repeats which were interspaced by only 1 bp into a unique telomeric repeat stretch by using the software mergeBed included in the suite bedtools (version 2.17.0) with the option “-d 1” (Quinlan & Hall, 2010). Then, we filtered the list to keep only the stretches longer than 10 bp. The stretches located at chromosome-ends were annotated as telomeres and were not kept for further analyses due to the unreliability of assembling softwares to correctly assemble chromosome-ends. The interstitial telomeric repeats stretches were annotated as Y’-associated ITS if they closely preceded an annotated Y’ element, while the others were discarded. The ITS content of a strain was determined as the sum of the lengths of its ITS annotations.

TL of these 12 yeast strains was determined by analysing their already available Sanger-sequencing reads, included in the SGRP project, as described in the paragraph “TL estimation in the SGRP collection”. Southern blots of 7 of these strains and of 17 representatives of the clades in the 918 yeasts collection were performed as previously described (Liti, Haricharan, et al., 2009).

### TL estimation in the SGRP collection

We estimated the TL of SGRP strains by using custom Perl scripts. First, we filtered the Sanger-sequencing reads to retain only the ones containing abundant telomeric repeats. We classified reads as telomeric if they contained a stretch of telomeric repeats (either C{1,3}A or TG{1,3}) longer or equal to 40 bp and located at the extremities of the read. Then, TL was estimated from each telomeric read by manually counting the telomeric repeats. Each read was considered as a single telomere.

### Statistics and data reproducibility

We estimated TL in 918 already sequenced *S. cerevisiae* strains (Peter et al., 2018). Strains classifications, including phylogenetic lineages and domesticated/wild assignments were maintained as described in (De Chiara, Barré, et al., 2020). We used the whole collection (*n*=918) for most of the analyses performed in this study, except for the GWAS (see below) and the correlation analysis (see paragraphs “Genome-wide association study and LOFs analysis” and “Correlation analysis”).

Statistical analyses were performed using R (version 3.5.3). Two-group comparisons were performed using the function *wilcox.test*, performing a two-tailed Mann-Whitney test. Multiple-group comparisons were performed using the function *kruskal.test*, performing a non-parametric ANOVA test. Subsequent pairwise analyses were performed using the function *dunnTest* included in the package FSA.

### Genome-wide association study and LOFs analysis

We used a subset of euploid diploid strains (*n*=555) for the genome-wide association study (GWAS) in order to minimize the confounding effects of ploidy variation and aneuploidy.

Genotype data and the matrix of between-strains co-similarity to adjust for population structure were already available from (Peter et al., 2018). Only variants with a minor allele frequency >= 0.05 in the whole population were included in the analysis (*n*=101579). TL data were converted to a normal distribution using the function *qqnorm* included in the R software. GWAS was performed using fast-lmm with the following command line:

~~~
fastlmmc -bfile $genotypes_file -bfileSim $cosimilarity_matrix -pheno $phenotypes_file -out $output_file
~~~

Variants whose corrected *p*<0.05 were considered statistically significant and retained for further analyses. Narrow-sense heritability (*h^2^*) for TL was calculated by dividing the genetic variance of the null model by the total variance of the null model (genetic variance and residual variance), computed using fast-lmm. The fraction of the phenotypic variance explained by the significant GWAS variants was calculated by removing all the rest of the genotypes from the co-similarity matrix and re-running a second GWAS. Subsequently, we repeated the same calculations on the new output file.

LOF variants were already available from (Peter et al., 2018). A LOF was considered as present in a strain if it was either in homozygous or heterozygous state. LOFs were annotated as present in a clade if their frequency in the clade was higher than 0.6 but their frequency in the 555 *S. cerevisiae* collection was lower than 0.2.

### Phenotypic measurements and correlation analysis

The set of phenotypes used in the correlation analysis was already available from previous studies. 35 phenotypes relative to resilience in the presence of stressors were derived from (Peter et al., 2018). 114 phenotypes relative to growth in multiple carbon and nitrogen sources were derived from (De Chiara, Barré, et al., 2020). Absolute and relative mtDNA copy number were derived from (De Chiara, Friedrich, et al., 2020). Mitochondrial volume and activity were partially derived from (De Chiara, Friedrich, et al., 2020) but, in this study, the measurements were extended to the whole 918 yeasts collection and were performed as described in (De Chiara, Friedrich, et al., 2020). However, only the euploid diploid strains (*n*=555) were used in the correlation analysis. Correlation analyses were performed using the function *cor.test* included in the R software and applying both Pearson’s and Spearman’s models. *P*-values were subsequently re-computed using both FDR and Bonferroni correction for multiple hypothesis testing and associations were considered statistically significant if their Spearman-derived FDR-corrected *p*<0.05.

## Data availability

The simulated reads of the synthetic genomes generated in this study are available upon request. The phenotype data and other resources are available at: https://github.com/mdangiolo89/Different-telomere-length-patterns-between-domestic-and-wild-S.-cerevisiae-isolates.

## Code availability

Y^ea^ISTY (Yeast ITS, Telomeres and Y’ elements estimator) is freely available at: https://github.com/mdangiolo89/Different-telomere-length-patterns-between-domestic-and-wild-S.-cerevisiae-isolates.

## Authors contributions

M.D., J.X.Y. and M.D.C. designed and implemented Y^ea^ISTY; M.D. and M.D.C. performed and analysed GWAS; B.B. and M.D. performed and analysed mitochondrial phenotyping; M.D. and M.J.G.P. performed teloblots; E.G. and G.L. conceived and supervised the project; M.D., E.G. and G.L. wrote the paper.

## Acknowledgements and funding

We thank the French National Research Agency (ANR) LABEX SIGNALIFE ANR-11-LABX-0028-01 and the Fondation pour la Recherche Médicale (FDT201904008453) for supporting M.D. PhD fellowship. This study was supported by ANR (ANR-18-CE12-0004, ANR-20-CE12-0020), Fondation pour la Recherche Médicale (EQU202003010413), CEFIPRA, Fondation ARC (N° PJA32020070002320) to G.L. Work in E.G. laboratory is supported by grants from the Fondation ARC pour la recherche sur le cancer (Labelisation N° PGA20160203873) and the INSERM cross cutting program on aging (AGEMED).

