## Supplementary text and figures for "Telomeres are shorter in wild *Saccharomyces cerevisiae* isolates than in domesticated ones"

### Supplementary discussion

#### 1, Implementing the Y$^{ea}$ISTY algorithm and benchmarking against simulated datasets

In this work, we developed a bioinformatic pipeline (Y$^{ea}$ISTY) that estimates telomere length (TL), ITS content and Y' elements copy number from whole-genome-sequencing data. The main idea at the basis of the Y$^{ea}$ISTY algorithm is that TL is directly proportional to the number of reads derived from telomeres. Therefore, correctly identifying telomeric reads is crucial to obtain reliable measurements. Yeast telomeric sequences are constituted by the repetition of very short motifs (TG1-3) that can also be found in internal parts of the genome. In addition, subtelomeric regions often harbor stretches of telomeric repeats (ITS) that can have lengths comparable to those of telomeres. We reasoned that the frequency distribution of the length of stretches of telomeric repeats in internal parts of the genome should be skewed towards smaller values than that of telomeric or subtelomeric regions, and this could constitute a first criterion to distinguish telomeric reads from the others. To test this, we derived the frequency distribution of the length of stretches of telomeric repeats in the complete genome assemblies of the yeast population reference panel (YPRP), comprising 7 *S. cerevisiae* and 5 *S. paradoxus* strains (Yue et al., 2017). In all the cases, the distribution was skewed towards small values, with an abundance of very short telomeric repeat stretches, mostly ranging between 2 and 20 bp. Very long stretches were exclusively present in subtelomeres (as ITS) or at telomeres (**Supplementary Fig. 1b and Supplementary Table 2**).

The next challenge was to find a suitable criterion to distinguish between telomeric and ITS reads. To do this, we explored the distribution of ITS and Y' elements in the YPRP. ITS content ranged from 0 to 3056 bp per haploid genome (median=549±1034 bp), while Y' copy number ranged from 0 to 31 (median=9.5±8.8). SK1 was particularly abundant in ITS/Y' (3056 bp, 31 copies), while N44 was totally depleted from them. We observed that subtelomeric ITS are always associated to a Y' element, while Y' elements can also exist without an associated ITS. Overall, ITS and Y' were positively correlated ($r$=0.7, $p$=0.01) (**Supplementary Fig. 1c and Supplementary Table 2**). Furthermore, tandemly-repeated sequences of ITS and Y' elements are occasionally observed in subtelomeric regions. We combined these two observations and reasoned that, since telomeres are the terminal parts of chromosomes while ITS are more internal, every read containing TG{1,3} repeats and whose pair maps on a Y' element must derive from a ITS. In contrast, reads containing C{1,3}A repeats whose pair maps on a Y' element cannot be unquestionably classified as ITS-derived, as they might derive either from a telomere or a tandemly-repeated ITS/Y' combination (**Supplementary Fig. 1a**).

Based on these observations, we tested the efficiency of Y$^{ea}$ISTY by using modified YPRP genome assemblies in which we simultaneously replaced all the native telomeres with 14 synthetic telomeres of known length, ranging from 18 to 651 bp, building a total of 168 synthetic genomes.

The use of the 12 YPRP backbone genomes, each one carrying variable ITS content, allowed us to test how much the genomic background and the variation in ITS content impact the TL estimation. We simulated Illumina paired-end reads from these genomes and applied $Y^{ea}$ISTY to these datasets using gradually increasing thresholds, ranging from 20 to 50 bp of telomeric repeats stretch, to check which one gave the best performance. The estimates of ITS content and TL correlated positively with the real values across the range of tested parameters, with the best correlation at the threshold of 40 bp ($r$=0.9842, $p$<2.2e$^{-16}$ for TL, $r$=0.9707, $p$<2.2e$^{-16}$ for ITS content) (**Supplementary Fig. 2a-b and Supplementary Table 3**). Furthermore, the Y' element set as additional chromosome entry in our modified reference genome enabled us to get an estimation of the copy number of these repetitive regions in our samples, by dividing the median Y' element coverage by the median coverage along the genome. We tested this method on our simulated datasets and found a significant positive correlation between our estimations and the values obtained from the annotation of the genome assemblies ($r$=0.89, $p$<2.2e$^{-16}$) (**Supplementary Fig. 2c)**. Overall, the results showed that $Y^{ea}$ISTY gives reliable estimations and the cutoff of 40 bp is the one giving the best results, therefore all the rest of the analyses shown in this study are performed using this threshold.

**2, Benchmarking of $Y^{ea}$ISTY against real experimental datasets**

We further validated the performance of $Y^{ea}$ISTY on real experimental datasets, by using the genome annotations of the YPRP, Sanger sequencing data derived from the *Saccharomyces* genome resequencing project (SGRP) and the standard technique of Southern blotting (teloblot) (Bergstrom et al., 2014; Liti et al., 2009; Yue et al., 2017). First, we retrieved the SGRP Sanger sequencing data corresponding to the 12 strains in the YPRP. We used custom Perl scripts to identify reads containing full-length telomeres and estimated TL by manually counting the telomeric repeats of those reads. In parallel, we estimated TL by applying $Y^{ea}$ISTY to Illumina sequencing data derived from the same strains (Yue et al., 2017).

We found significant positive correlations for TL, ITS content and Y' copy number (Pearson's $r$=0.63 and $p$=0.04, $r$=0.89 and $p$=9.63e$^{-5}$, $r$=0.83 and $p$=0.0008, respectively) (**Supplementary Fig. 3a and Supplementary Table 4**). In addition, we measured TL of 5 *S. cerevisiae* strains (DBVPG6044, DBVPG6765, Y12, YPS128, UWOPS034614) and 2 *S. paradoxus* strains (CBS432 and N44) by teloblot and confirmed a positive correlation between all the measurements (Pearson's $r$=0.99 and $p$=0.0001 between the Sanger sequencing and the teloblot, $r$=0.78 and $p$=0.06 between the teloblot and $Y^{ea}$ISTY). One strain, N44, did not give any terminal restriction fragment (TRF) due to the complete absence of Y' elements in its genome and was not included in the statistical analysis (**Supplementary Fig. 3b-d and Supplementary Table 4**). We observed underestimation for both TL and ITS content as compared to the values measured by teloblot and the annotation of the 12 YPRP genome assemblies. The underestimation rate for TL in this experimental setting was higher than in the 918 *S. cerevisiae* collection (9.6 vs 4-fold) and it implies that the sequencing technology has an impact on the estimation and samples derived from different sequencing batches cannot be directly compared. Nevertheless, the data show that Y$^{ea}$ISTY performs well also on real experimental datasets and can be used for further analyses.

**3, Application of Y$^{ea}$ISTY to multiple experimental settings**

The recent publication of the complete genome sequencing of the *Saccharomyces cerevisiae* knock-out collection and its use to identify genes involved in genome stability, including TLM ones, offered the opportunity to test Y$^{ea}$ISTY against an already validated dataset (Puddu et al., 2019). Puddu *et al.* applied a bioinformatic approach that is similar to the one used in our study, and identified, among others, 12 new TLM genes validated by teloblot. We applied Y$^{ea}$ISTY to the same dataset, adding knock-outs for known telomere-associated proteins, like *RIF1*, *RIF2*, *TEL1* and the subunits of the telomerase complex (*EST1*, *EST2*, *EST3*). In all the cases Y$^{ea}$ISTY detected a significant difference in TL respect to the wild-type samples and the results were in line with what previously described in Puddu *et al*. (Pearson's $r$=0.86, $p$=1.38e$^{-15}$), confirming the reliability of our approach. In only one case (*UAF30Δ*), our predictions were in contrast with those of Puddu *et al.* In the case of *EST* genes, we found that their high TL estimation partly derives from amplification of ITS/Y' elements **(Supplementary Fig. 4a-d and Supplementary Table 4)**.

Next, we tested whether Y$^{ea}$ISTY was able to detect changes in TL, ITS content and Y' copy number across a long-term experimental evolution by re-analysing diploid *S. cerevisiae* and *S. paradoxus* strains and their hybrids (both intra- and inter-specific), which had been evolved through a mutation accumulation lines (MALs) protocol for 120 single-cell bottlenecks (scb, ~2400 generations) (Tattini et al., 2019). Our analyses revealed that, while the majority of the evolved strains maintained their TL, ITS content and Y' copy number constant over time, the diploid *S. paradoxus* (N17xN17) evolved strains underwent massive telomere elongation, accompanied by ITS/Y' amplification, and resembled type I survivors. On the contrary, their founder strain had TL and ITS/Y' comparable to the other genetic backgrounds (**Supplementary Fig. 4e and Supplementary Table 4)**. Given that all the *S. paradoxus* lines underwent telomere elongation and we could not identify the cause in any of the new mutations occurred during the evolution protocol, we conclude that the phenotype must derive from the genetic background of their founder strain, which might contain variants that promote Y' recombination and expansion upon continuous propagation. In accordance with this hypothesis, their *S. paradoxus* founder strain (N17) belongs to the European *S.*

*paradoxus* population, which is known to harbour very short telomeres that might be the result of mutations in TLM genes (Liti et al., 2009).

Taken together, these results show that Y$^{ea}$ISTY can be applied to multiple experimental settings and can unravel telomeric and subtelomeric modifications.

**4, Telomere length, ITS content and Y' copy number are closely associated**

We measured ITS content across the 918 *S. cerevisiae* collection and found it to have a hyperbolic distribution, with the majority of the strains containing less than 1000 bp of ITS per haploid genome. The overall distribution ranged from 0 to 74312 bp per haploid genome (median=1036±4541 bp) (**Supplementary Fig. 6a and Supplementary Table 5**). We subsequently estimated ITS content in the clades and found it to vary significantly among them (two-tailed Kruskal-Wallis test, $p<2.2e^{-16}$). It was highest in Mexican agave and Brazilian bioethanol, where the median exceeded 4000 bp per haploid genome, while it was lowest in Mediterranean oak, Ecuadorean and CHNV clades, where ITS were almost absent (**Supplementary Fig. 6b and Supplementary Table 7**). ITS distribution in the Wine European clade was similar to the whole collection and ITS content varied significantly among the Wine European subclades (two-tailed Kruskal-Wallis test, $p=8.95e^{-14}$). It followed the same trend of TL, being highest in the Clinical/Y' amplification subclade where the median exceeded 40000 bp, and lowest in the Clinical/*S. boulardii* subclade, where it remained below 100 bp (**Supplementary Fig. 6c-d and Supplementary Table 7**).

In yeast, subtelomeric ITS are usually associated to Y' elements. Therefore, we estimated the Y' copy number in the 918 *S. cerevisiae* collection to see if it correlated with ITS content. Y' copy number had a distribution similar to ITS content, with the majority of the strains containing ~15 Y' elements (**Supplementary Fig. 7a and Supplementary Table 5**). Its variation was wide across the lineages (two-tailed Kruskal-Wallis test, $p<2.2e^{-16}$) and followed the ITS content distribution for most of the clades. It was highest in Mexican agave, Mixed origin, Brazilian bioethanol, Asian islands and Ale beer, all containing more than 20 Y' elements. It was lowest in the Ecuadorean, CHNV, Far East Asia, CHNIII and Mediterranean oak, all containing less than 3 Y' elements. The Ecuadorean clade, in particular, contained no Y' elements at all (**Supplementary Fig. 7b and Supplementary Table 8**). Interesting exceptions are the French dairy, African beer, Ale beer, African palm wine and Asian islands clades, which contain very few ITS and a high number of Y' elements. The opposite case, with high ITS and few Y' elements, was never found (**Supplementary Fig. 6b and 7b**). Y' copy number also varied among the subclades of the Wine European clade (two-tailed Kruskal-Wallis test, $p<2.2e^{-16}$) and it followed the same trend of ITS content, being highest in the Clinical/Y' amplification subclade, where the median exceeded 100 Y' elements, and lowest in the Clinical/*S. boulardii* subclade, where it remained below 5 copies (**Supplementary Fig. 7c-d)**.

The similar behaviour of TL, ITS content and Y' copy number suggests that they might be closely associated. We measured the correlation coefficients among the three variables and found significant positive correlations (Pearson's $r$=0.43, $p$<2.2e$^{-16}$ between TL and ITS, $r$=0.81 and $p$<2.2e$^{-16}$ between Y' and ITS, $r$=0.39 and $p$<2.2e$^{-16}$ between TL and Y') (**Supplementary Fig. 8a**). This result suggests that either ITS/Y' might be *cis*-acting factors influencing the homeostasis of TL or viceversa (Brevet et al. 2003; Craven & Petes, 1999), or that TL, ITS content and Y' copy number are under the control of the same factors. To test these hypotheses, we measured TL, ITS content and Y' copy number in knock-out strains for known TLM genes (Puddu et al., 2019) and detected a significant difference in both TL and ITS/Y' respect to the wild-type samples. In most of the cases, the direction of the variation for ITS and Y' was the same as for TL, although some exceptions were also observed (*AIM4Δ, SLM4Δ, RIF1Δ* for ITS content, *AIM4Δ, MDY2Δ* for Y' copy number). This confirms that ITS, Y' and telomeres are positively correlated (Pearson's $r$=0.65 and $p$=4.841e$^{-7}$ between TL and ITS, $r$=0.69 and $p$=3.99e$^{-8}$ between TL and Y', $r$=0.96 and $p$<2.2e$^{-16}$ between ITS and Y') and are under the control of the same genetic determinants (**Supplementary Fig. 4b-d and Supplementary Table 4**).

We then checked how ITS and Y' behaved in domesticated and wild yeasts. Both ITS content and Y' copy number were lower in wild yeasts than in domesticated ones (two-tailed Wilcoxon test, $p$=1.35e$^{-5}$ and $p$<2.2e$^{-16}$ for ITS and Y', respectively) (**Supplementary Fig. 8b-c**). Among the wild clades, the Taiwanese, Russian and North-American behaved like domesticated ones, having high ITS content and Y' copy number (**Supplementary Fig. 6b and 7b**). We investigated whether domestication played a role in the association between TL and ITS content. However, TL and ITS/Y' were always correlated to each other even if we considered only domesticated (Pearson's $r$=0.48 and $p$<2.2e$^{-16}$ between TL and ITS, $r$=0.44 and $p$<2.2e$^{-16}$ between TL and Y', $r$=0.84 and $p$<2.2e$^{-16}$ between ITS and Y') or only wild isolates (Pearson's $r$=0.36, $p$=0.007 between TL and ITS, $r$=0.35 and $p$=0.007 between TL and Y', $r$=0.92 and $p$<2.2e$^{-16}$ between ITS and Y'), meaning that this association is mediated by other factors. No variation in ITS content was detected between domesticated-anthropic and domesticated-feral strains (two-tailed Wilcoxon test, $p$=0.49) while the two groups differed significantly in terms of their Y' copy number (two-tailed Wilcoxon test, $p$=0.0009) (**Supplementary Fig. 8b-c**).

In conclusion, we show that telomeres and ITS/Y' biology are closely associated and interdependent, and that telomeres and ITS/Y' are under the control of the same genetic factors.

**5, Loss-of-function variants in TLM genes potentially contributing to TL variation**

We checked if the presence of rare (frequency<20% in the 555 *S. cerevisiae* collection) loss-of-function variants (LOFs) could explain extreme TL patterns observed in some clades (**Supplementary Table 11**). Among the ones with the shortest telomeres, the Malaysian clade carries multiple chromosomal rearrangements (Yue et al., 2017), which coincide with a fixed LOF in the gene *SIR4*, whose deletion causes telomere shortening and genome instability, or in *HUR1*, a gene involved in non-homologous end-joining repair. A LOF in *URN1*, causing telomere shortening, was detected in the Ecuadorean strains. The clinical/*S. boulardii* subclade carried LOFs in *AZR1* and *WHI2*, whose deletion leads to telomere shortening. On the contrary, a third LOF in the gene *RPS30B* causes telomere lengthening. A LOF in *RNR1*, encoding a subunit of the ribonucleotide-diphosphate reductase, is fixed in the North American clade. The inactivation of this gene causes telomere shortening due to reduced availability of dNTPs and is in accordance with the phenotype of the clade. Among the lineages with the longest telomeres, the clinical/Y' amplification, Alpechin and Brazilian bioethanol did not carry any LOF. However, the *RIF2* gene of all the Alpechin strains is introgressed from *S. paradoxus* and we found a significant positive correlation (Pearson's $r$=0.82 and $p$=0.0003) between TL and the amount of introgressed genome (D'Angiolo et al., 2020). The French dairy clade contained three LOF variants (*HUR1, MND2, GUP2*), including one in the gene *GUP2*, whose deletion causes telomere lengthening. The African beer clade presented LOF variants in both telomere lengthening (*YBR284W*) and shortening (*MET7*) genes.

Overall, LOF variants in TLM genes that are present in specific clades or isolates might have large effect size contribution to the global TL variation, but they escape GWAS detection due to their low frequencies. Reverse engineering of these variants is required to estimate their effect and their interaction with the genetic background (De Chiara et al., 2020).

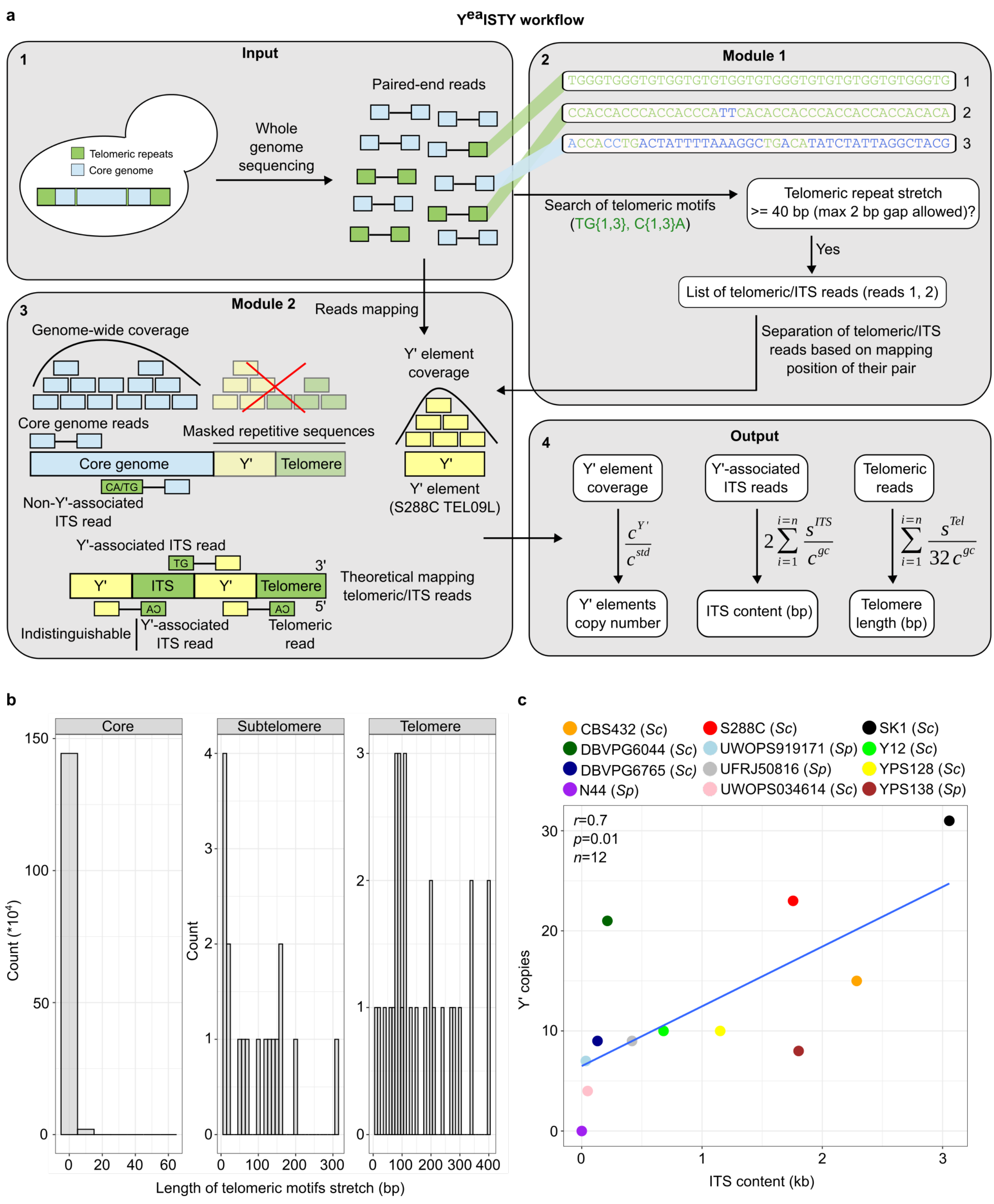


**Supplementary Figure 1 – Implementation of the Y$^{ea}$ISTY algorithm. a**, Paired-end reads are scanned to find telomeric sequence motifs (either C{1,3}A or TG{1,3}, in green). Reads containing a stretch of telomeric motifs longer or equal to 40 bp, possibly interspaced by one gap of maximum 2 nt, are retained (reads 1 and 2, but not 3) (panels 1, 2). All the reads (retained and non-retained) are mapped to a modified reference genome in which all repetitive sequences, including telomeres (in pale green) and Y' elements (in pale yellow), have been masked. Moreover, a long-version (6.7 kb) Y' element from the left telomere of chromosome IX of the lab strain S288C has been appended to this modified reference genome as an additional chromosome entry. The mapping position of retained reads and their pairs is investigated to classify them as: non-Y'-associated ITS reads if they contain a stretch of either C{1,3}A or TG{1,3} motifs and their pair maps on a core part of the genome (in light blue); Y'-associated ITS reads if they contain a stretch of TG{1,3} motifs and their pair maps on the appended Y' element; all the reads not fulfilling either of these criteria are classified as telomeric. For clarification of this principle, the figure shows an example of chromosome-end harbouring multiple tandemly-repeated Y' element/ITS combinations. Reads containing C{1,3}A motifs whose pair maps on the Y' element can equally derive from telomeres or Y'-associated ITS, whereas reads containing TG{1,3} motifs whose pair maps on the Y' element unambiguously derive from telomeres (panel 3). The amount of reads in each category is then used to infer ITS content and telomere length, while coverage data on the appended Y' element is compared to the genome-wide coverage to infer the copy number of Y' elements (panel 4). **b**, Example of frequency distribution of stretches of C{1,3}A and TG{1,3} repeats in the genome assembly of the *S. cerevisiae* laboratory strain S288C. Bin width is 10 bp. **c**, Comparison of ITS content (x axis) and Y' elements copy number (y axis) for the 12 *S. cerevisiae* (*Sc*) and *S. paradoxus* (*Sp*) strains in the YPRP.

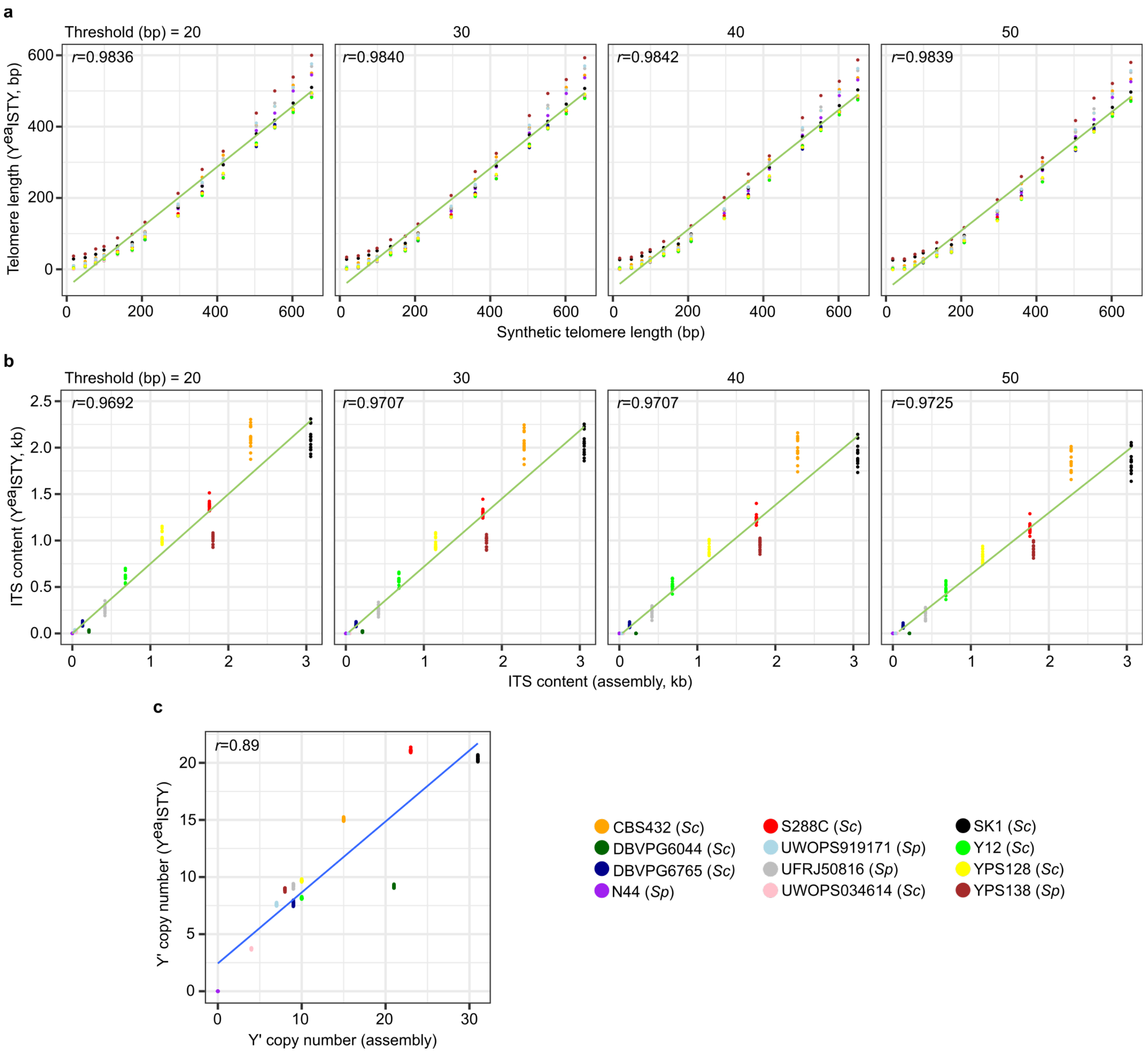


**Supplementary Figure 2 – Validation of Y$^{ea}$ISTY against simulated datasets. a-c**, The panels show estimated (y axis) and real (x axis) TL/ITS/Y' in 168 simulated sequencing runs with a coverage of 30X based on the genome assemblies in the YPRP, modified to carry synthetic telomeres of increasing length. Each point represents a single simulated Illumina paired-end sequencing run and the line represents a linear regression function. Side-by-side panels represent correlations obtained by running Y$^{ea}$ISTY with increasing thresholds of telomeric repeat stretches (20 to 50 bp) to detect telomeric reads. Y' copy number estimation is not affected by the chosen threshold, therefore only one panel is shown. $p$<2.2e$^{-16}$ and $n$=168 for all the panels.

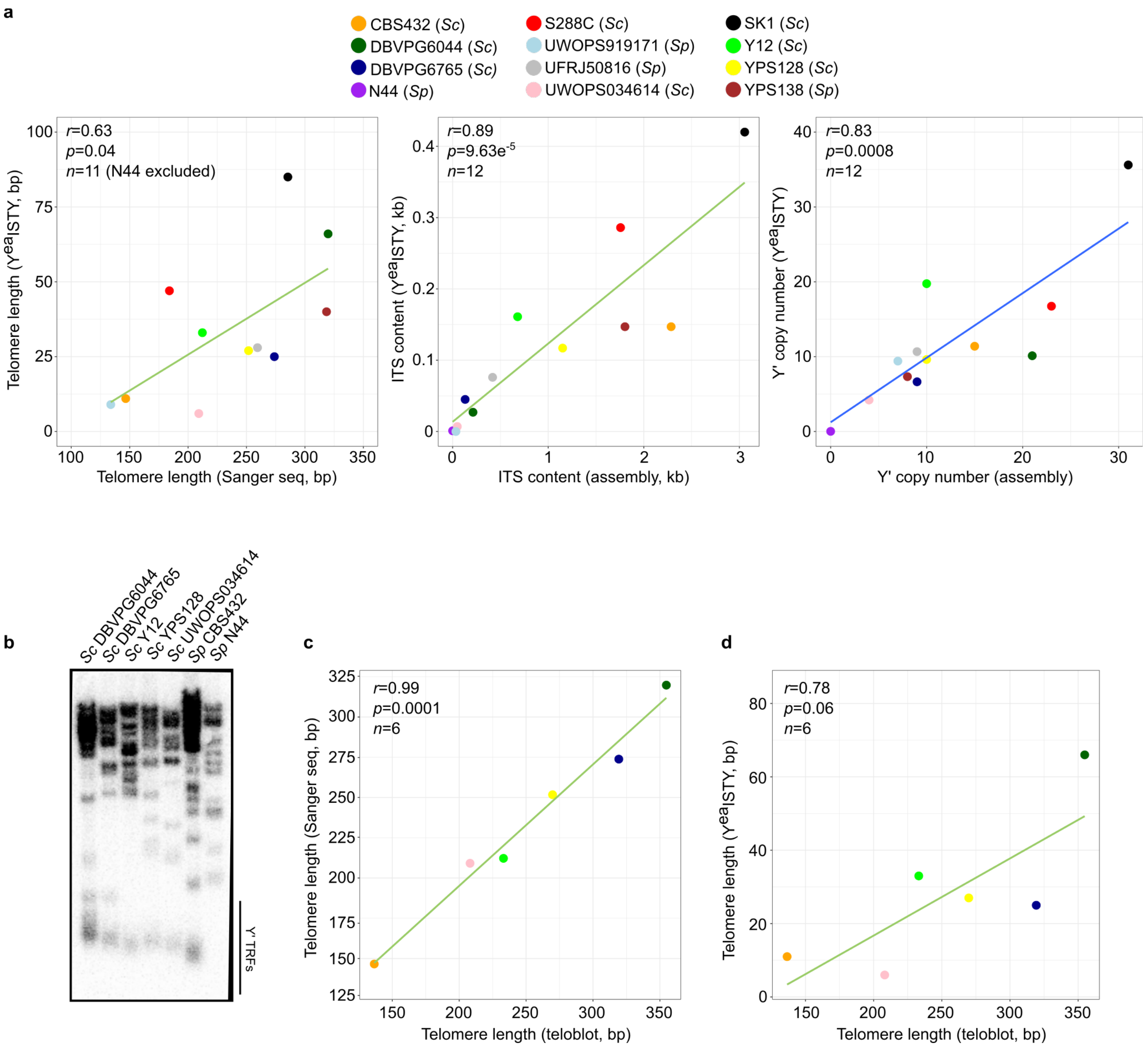


**Supplementary Figure 3 – Validation of Y$^{ea}$ISTY against real datasets. a**, Comparison of telomere length, ITS content and Y' copy number in the 12 strains of the YPRP, estimated by Y$^{ea}$ISTY (y axis) vs Sanger sequencing (for telomere length, left panel) or from the annotation of the genome assemblies (for ITS content and Y' copy number, middle and right panels) (x axis). **b**, *XhoI* digestion of genomic DNA derived from a subset of the YPRP strains. Genomic DNA is probed with radioactively-labelled telomeric TG1–3 repeats. The black line denotes terminal restriction fragments (TRFs) resulting from the digestion of a Y' element. No TRF was detected for the strain N44 because it does not carry any Y' elements. **c**, Comparison of telomere length in the same subset of strains, estimated by Sanger sequencing (y axis) and the teloblot (x axis). **d**, Comparison of telomere length in the same subset of strains, estimated by Y$^{ea}$ISTY (y axis) and the teloblot (x axis). Each point represents a single isolate and the line represents a linear regression function. Colours are as in plot **a**.

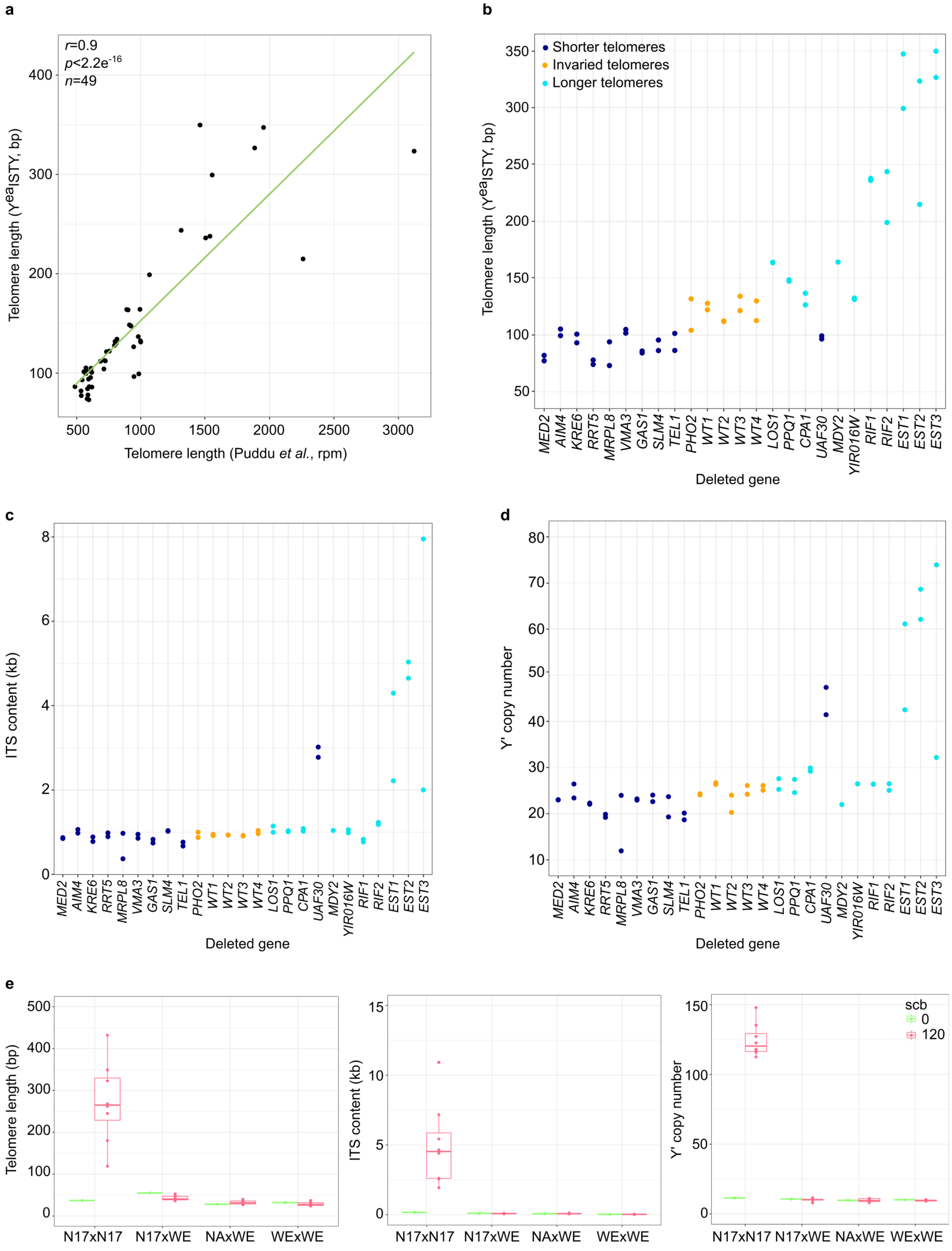


**Supplementary Figure 4 – Application of Y$^{ea}$ISTY to multiple experimental setups**. **a**, Comparison of TL estimations from Y$^{ea}$ISTY (y axis) vs those from Puddu *et al*. (x axis) (left). The line represents a linear regression function. rpm=telomeric reads per million. **b-d**, TL/ITS/Y' of knocked-out strains for known telomere length maintenance genes described in Puddu *et al*. (x axis), estimated by Y$^{ea}$ISTY (y axis). Colours are as in Puddu *et al.* and denote either wild-type strains or a non-TLM gene (*PHO2*, orange), or gene knock-outs that shorten (dark blue) or lengthen (light blue) telomeres. **e**, Estimation of TL, ITS content and Y' copy number before (green) and after (pink) an experimental evolution protocol, in crosses derived from Tattini *et al.* and evolved through 120 single-cell bottlenecks (scb). N17=*S. paradoxus*, WE=Wine European *S. cerevisiae*, NA=North American *S. cerevisiae*. *S. paradoxus* diploid strains (N17xN17) elongate telomeres and undergo massive amplification of ITS and Y' elements.

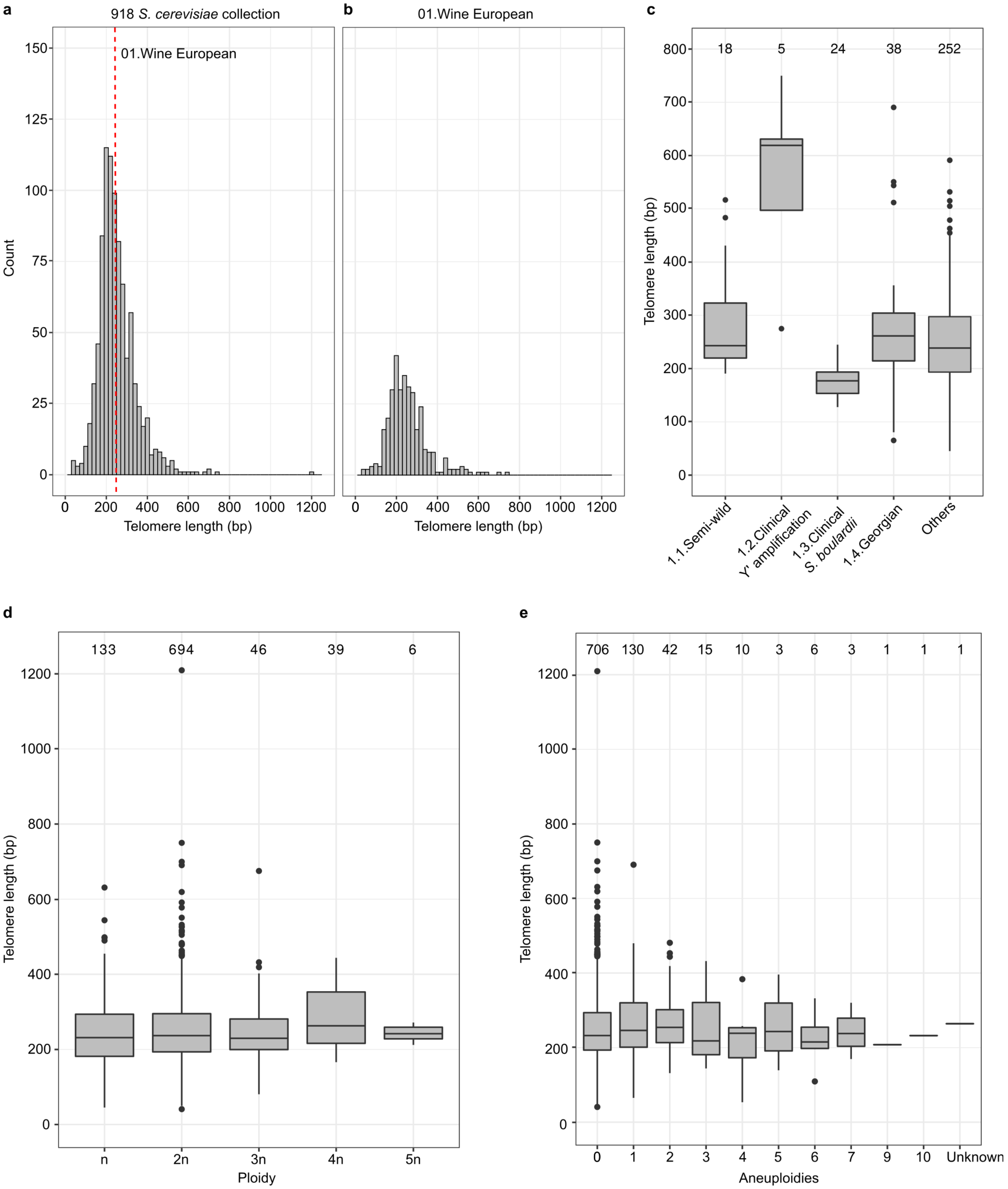


**Supplementary Figure 5 – TL variation in the 918 *S. cerevisiae* collection**. **a**, Frequency distribution of telomere length in the 918 *S. cerevisiae* collection. Bin width is 20 bp. The red line indicates the median of the Wine European clade. **b**, Frequency distribution of telomere length in the Wine European clade. Bin width is 20 bp. **c**, Telomere length in the subclades of the Wine European lineage. **d-e**, Telomere length of isolates grouped by their ploidy (panel **d**) or by their number of aneuploidies (panel **e**). Box plots are as in Fig. 2. Numbers on top represent the number of isolates in each group.

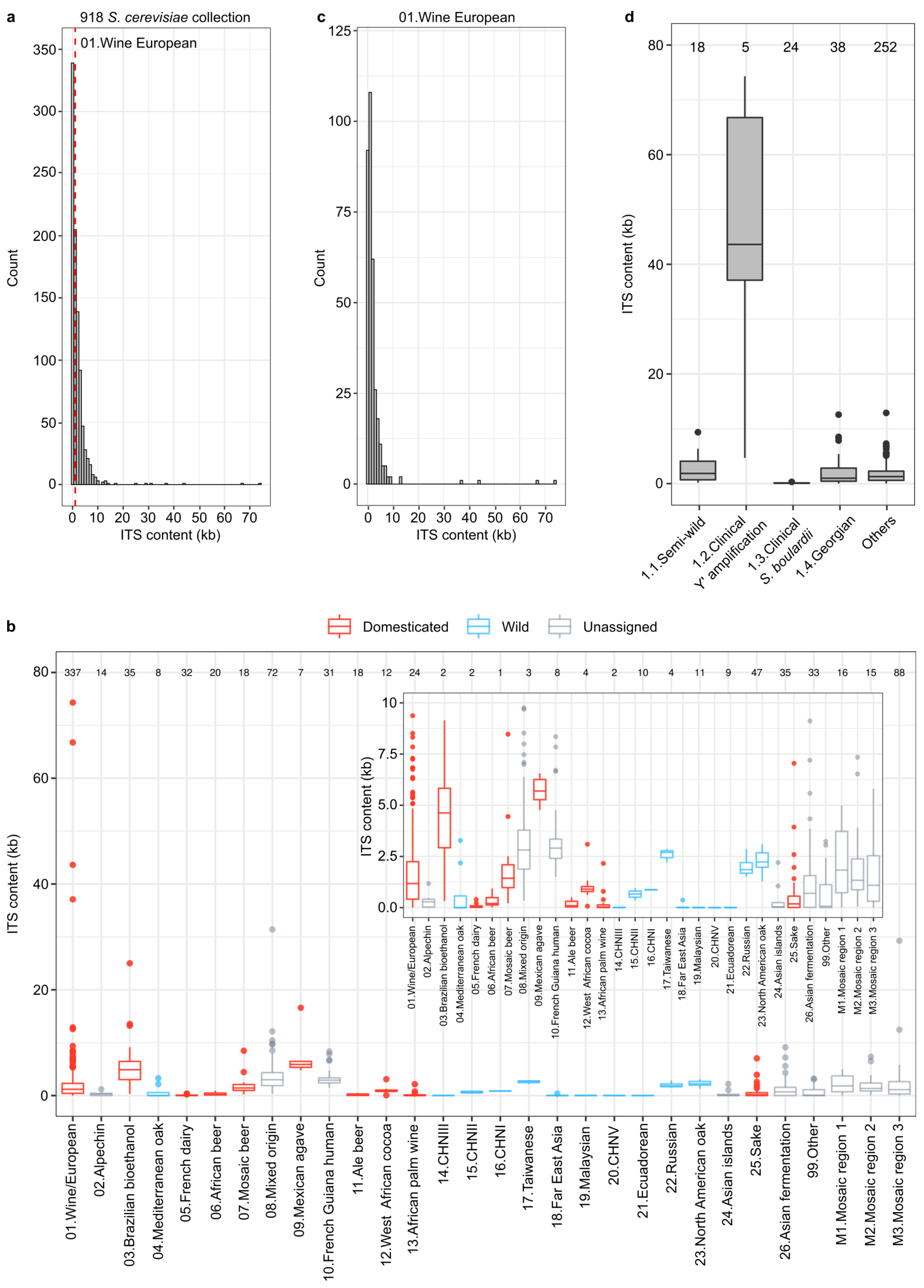


**Supplementary Figure 6 – ITS content variation in the 918 *S. cerevisiae* collection**. **a**, Frequency distribution of ITS content in the 918 *S. cerevisiae* collection. Bin width is 1 kb. The red line indicates the median of the Wine European clade. **b**, ITS content of the lineages in the 918 *S. cerevisiae* collection. Colours represent the clade classification (domesticated, wild or unassigned), which is the same as in Fig. 2. The order of clades on the x axis is as in Fig. 2. The inset shows a magnified view of the y axis up to 10 kb. **c**, Frequency distribution of ITS content in the Wine European clade. Bin width is 1 kb. **d**, ITS content in the subclades of the Wine European lineage. Boxplots are as in Fig. 2. Numbers on top represent the number of isolates in each subclade.

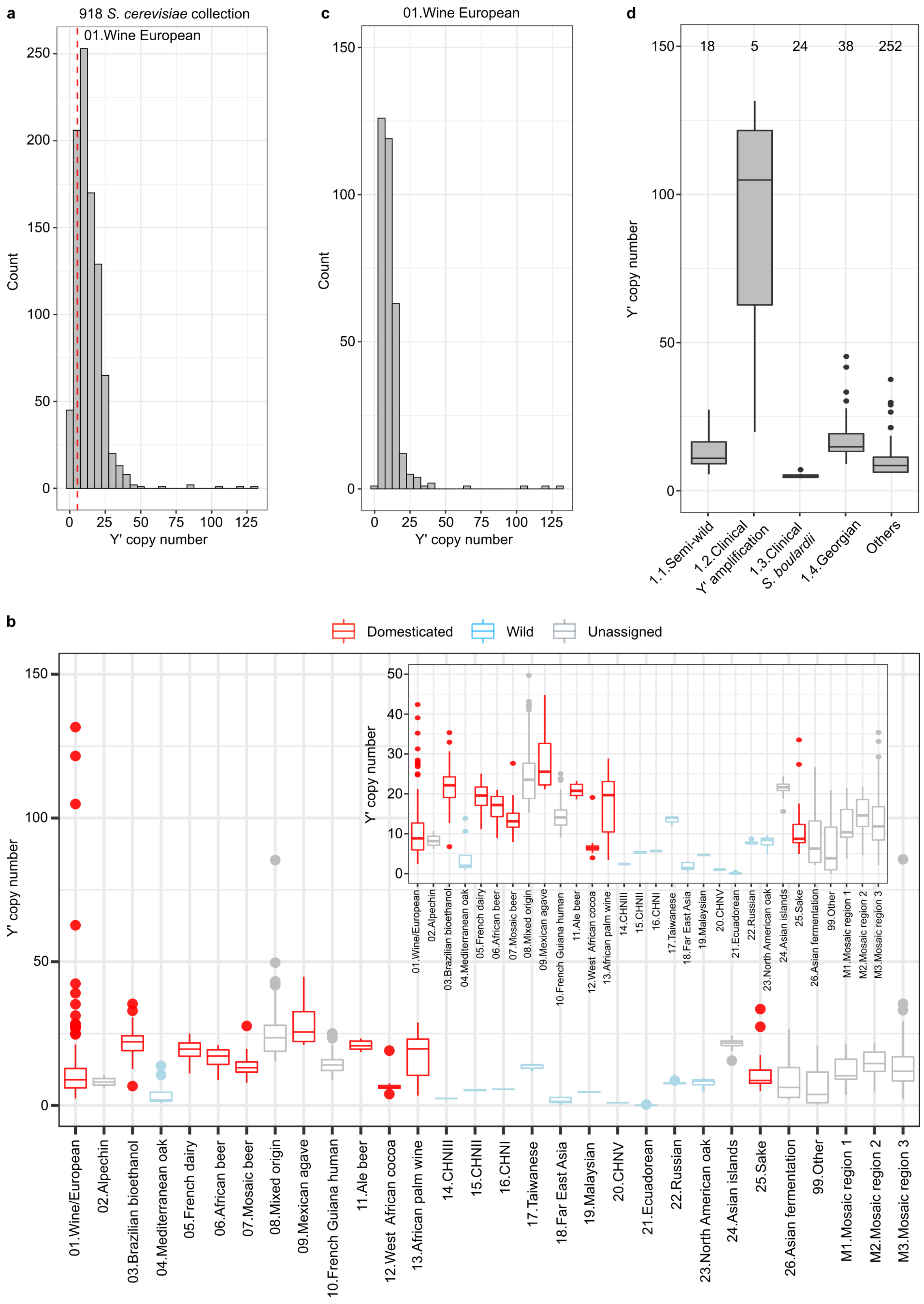


**Supplementary Figure 7 – Y' copy number variation in the 918 *S. cerevisiae* collection**. **a**, Frequency distribution of Y' copy number in the 918 *S. cerevisiae* collection. Bin width is 5 copies. The red line indicates the median of the Wine European clade. **b**, Y' copy number of the lineages in the 918 *S. cerevisiae* collection. Colours represent the clade classification (domesticated, wild or unassigned), which is the same as in Fig. 2. The order of clades on the x axis is as in Fig. 2. The inset shows a magnified view of the y axis up to 50 copies. **c**, Frequency distribution of Y' copy number in the Wine European clade. Bin width is 5 copies. **d**, Y' copy number in the subclades of the Wine European lineage. Boxplots are as in Fig. 2. Numbers on top represent the number of isolates in each subclade.

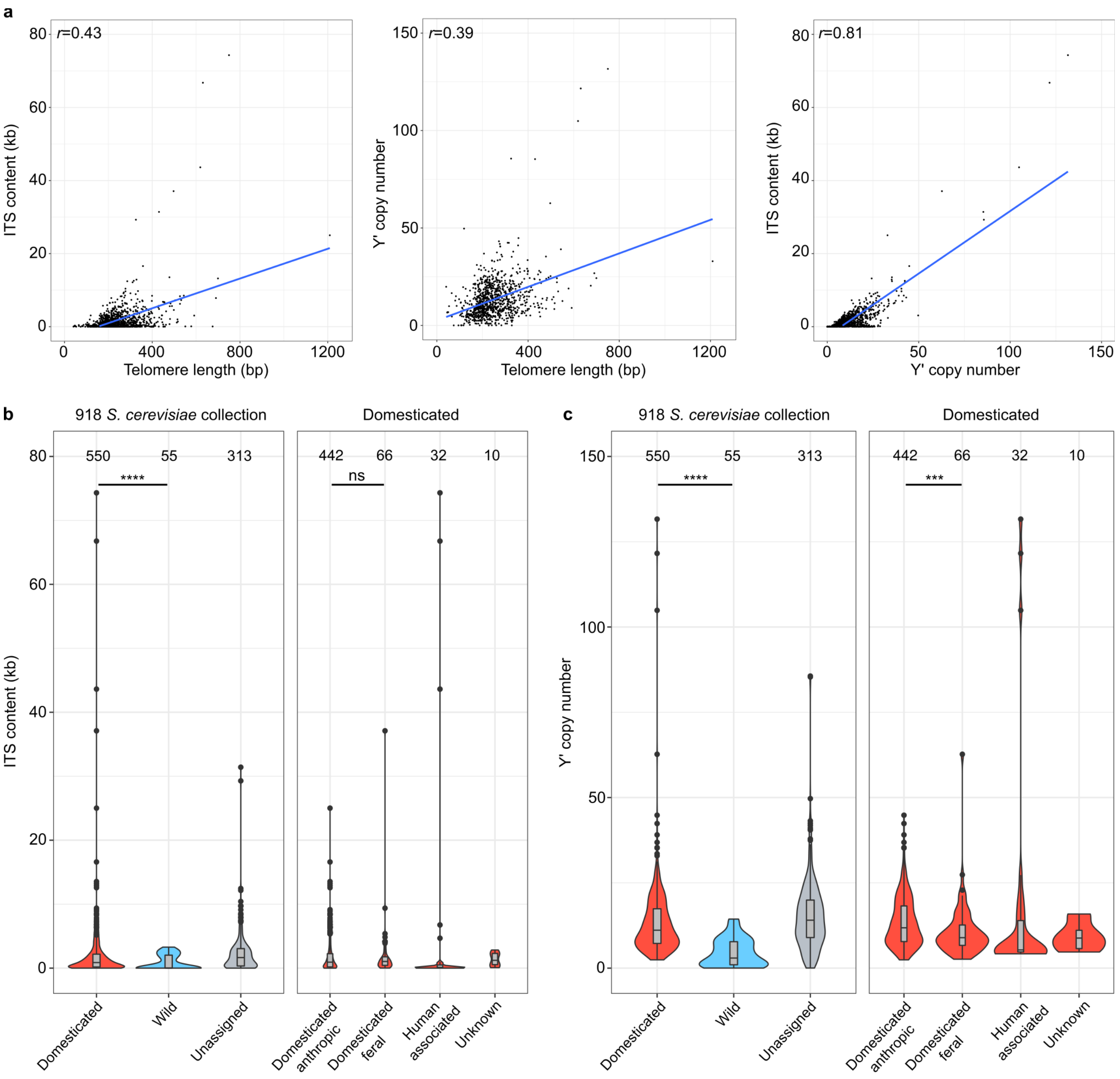


**Supplementary Figure 8 – Relationships among TL, ITS content and Y' copy number. a**, Correlation between TL and ITS content (left panel), TL and Y' copy number (middle panel), ITS content and Y' copy number (right panel) in the 918 *S. cerevisiae* collection. Each point represents a single isolate and the line represents a linear regression function. $p<2.2e^{-16}$ and $n$=918 for all the panels. **b-c**, ITS content (panel **b**) and Y' copy number (panel **c**) of domesticated vs wild isolates in the 918 *S. cerevisiae* collection (left panels), or of anthropic vs feral isolates in the domesticated isolates of the same collection (right panels). Box plots are as in Fig. 2. Numbers on top represent the number of isolates in each group. *$p < 0.05$, **$p < 0.01$, ***$p < 0.001$, ****$p < 0.0001$, ns=non-significant.

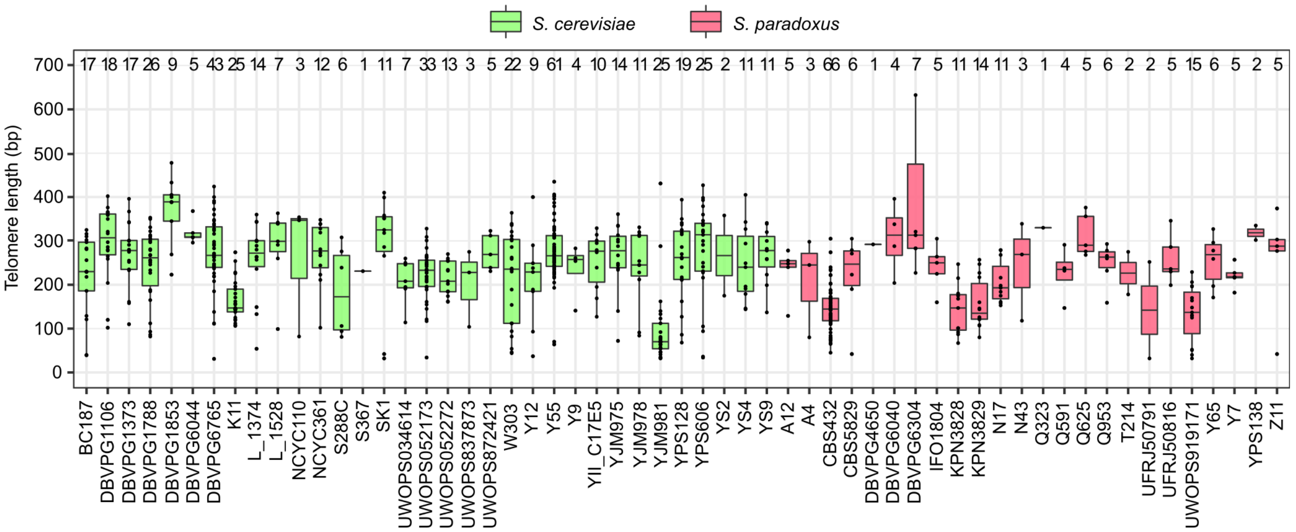


**Supplementary Figure 9 – TL variation in the SGRP collection.** Telomere length of strains in the *Saccharomyces* genome resequencing project (SGRP). Colours represent *S. cerevisiae* (green) and *S. paradoxus* (pink) isolates. Box plots are as in Fig. 2. Numbers on top represent the number of telomeric reads used to estimate the telomere length in each isolate.

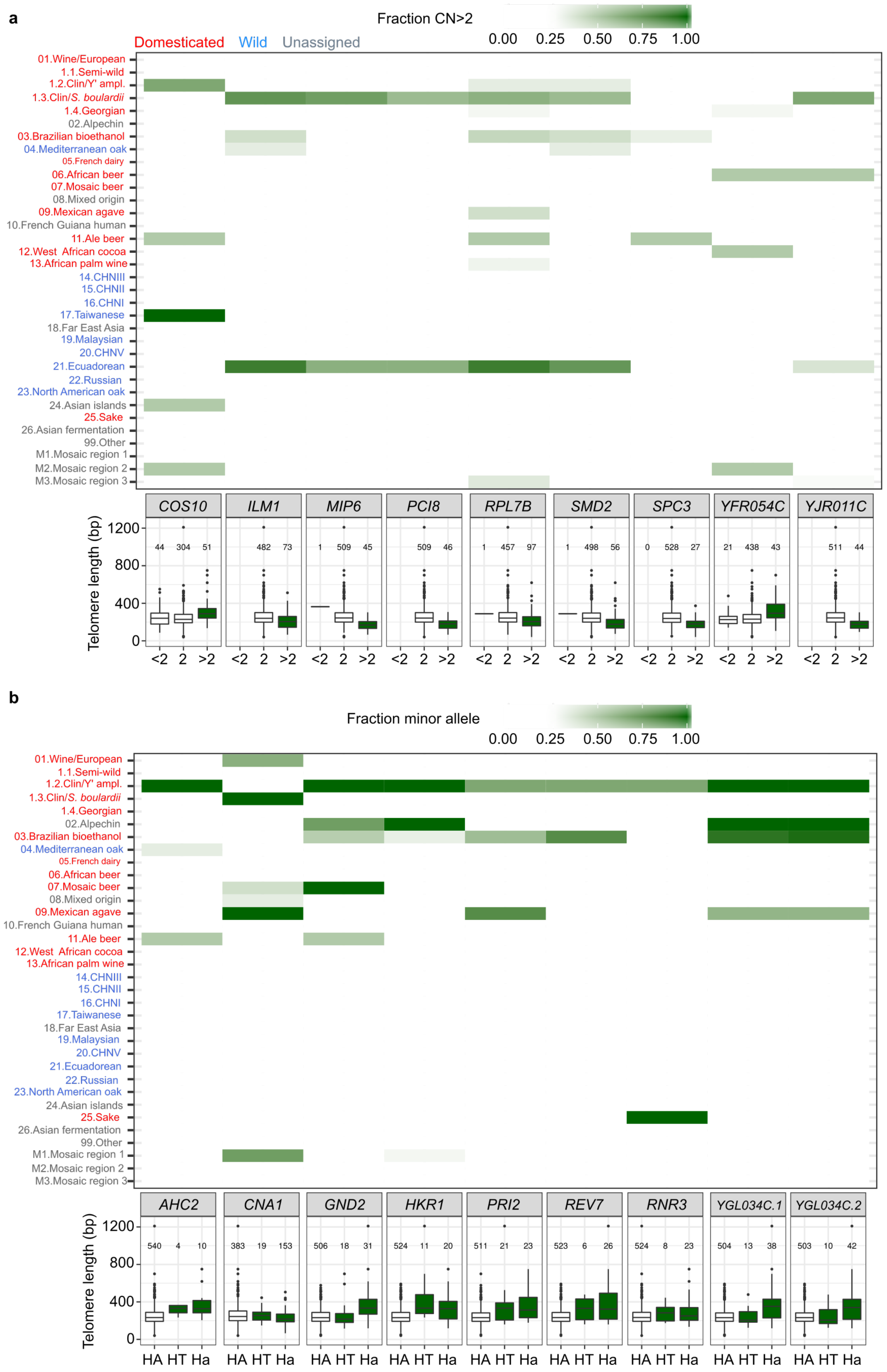


**Supplementary Figure 10 – Frequency and effect of GWAS variants. a,** The upper plot represents a heatmap of the gene copy number (CN) of the 9 significant copy number variants in the nuclear genome. Squares colour represents the fraction of isolates carrying more than 2 copies of the gene in each clade, and the intensity of the green tone is directly proportional to the fraction in the clade. The bottom plot represents the telomere length of isolates grouped by their CN for each of the variants. <2: less than 2 copies of the gene; 2: 2 copies of the gene; >2: more than 2 copies of the gene. The isolates whose gene CN was unknown are not displayed. **b**, The upper plot represents a heatmap of the minor allele frequency (MAF) of the 9 significant single nucleotide variants. Squares colour represents the MAF in each clade, and the intensity of the green tone is directly proportional to the MAF in the clade. The minor allele is considered as present in an isolate if it is either in homozygous or in heterozygous state. The order and colour of the clades on the y axis is the same as in Fig. 2. The bottom plot represents the telomere length of isolates grouped by their genotype. HA=homozygous for the major allele, HT=heterozygous, Ha: homozygous for the minor allele. Numbers on top represent the number of isolates in each group. The isolates whose genotype was unknown are not displayed. Box plots are as in Fig. 2.

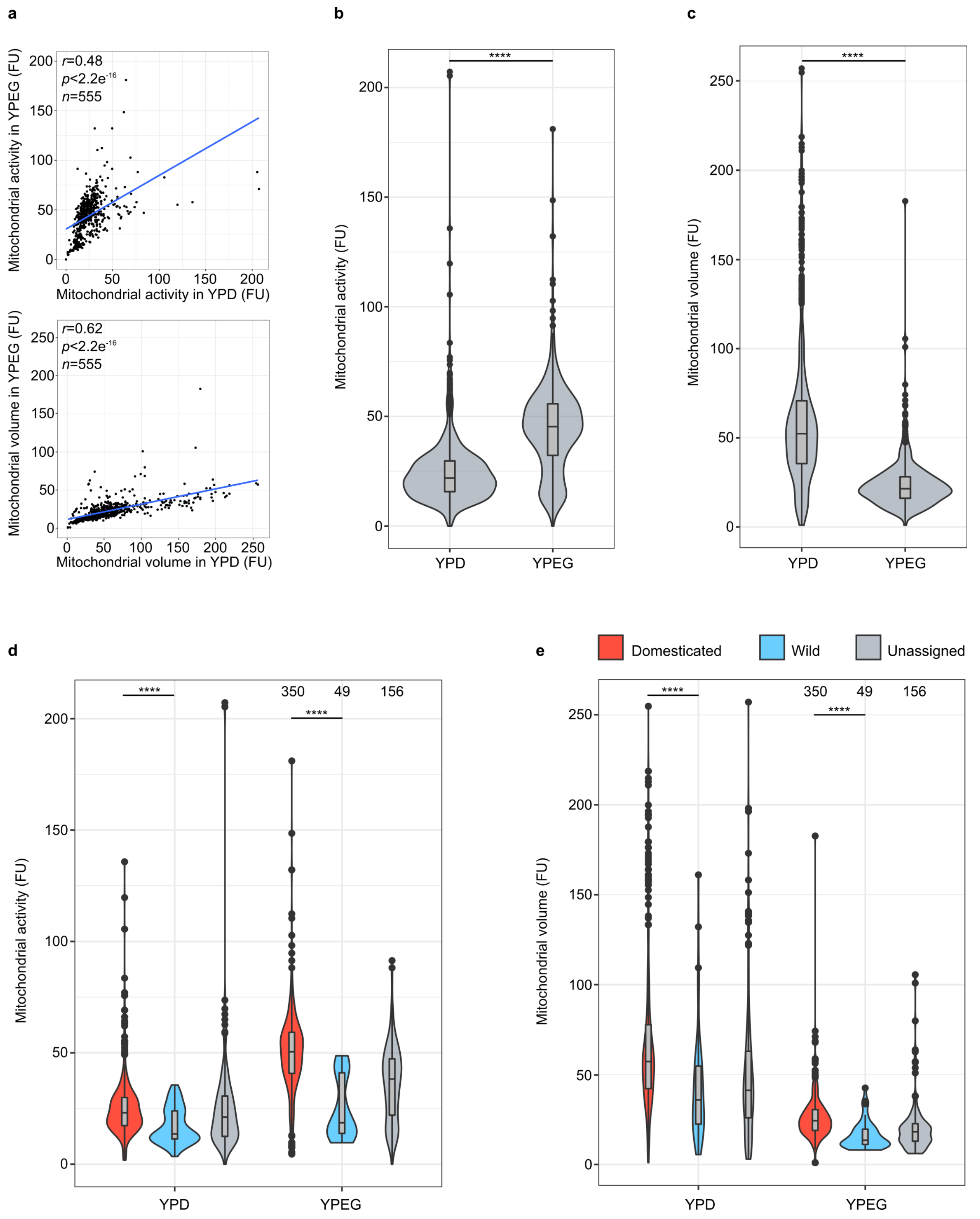


**Supplementary Figure 11 – TL and mitochondrial traits. a,** Correlation between mitochondrial activity (top) and volume (bottom) in YPD and YPEG media. Each point represents a single isolate and the line represents a linear regression function. FU=fluorescence units. **b-c,** Mitochondrial activity and volume in YPD and YPEG in the 555 *S. cerevisiae* collection. **d-e,** Mitochondrial activity and volume in YPD/YPEG of domesticated, wild and unassigned isolates in the 555 *S. cerevisiae* collection. Box plots are as in Fig. 2. Numbers on top represent the number of isolates in each group. *$p$ < 0.05, **$p$ < 0.01, ***$p$ < 0.001, ****$p$ < 0.0001, ns=non-significant.

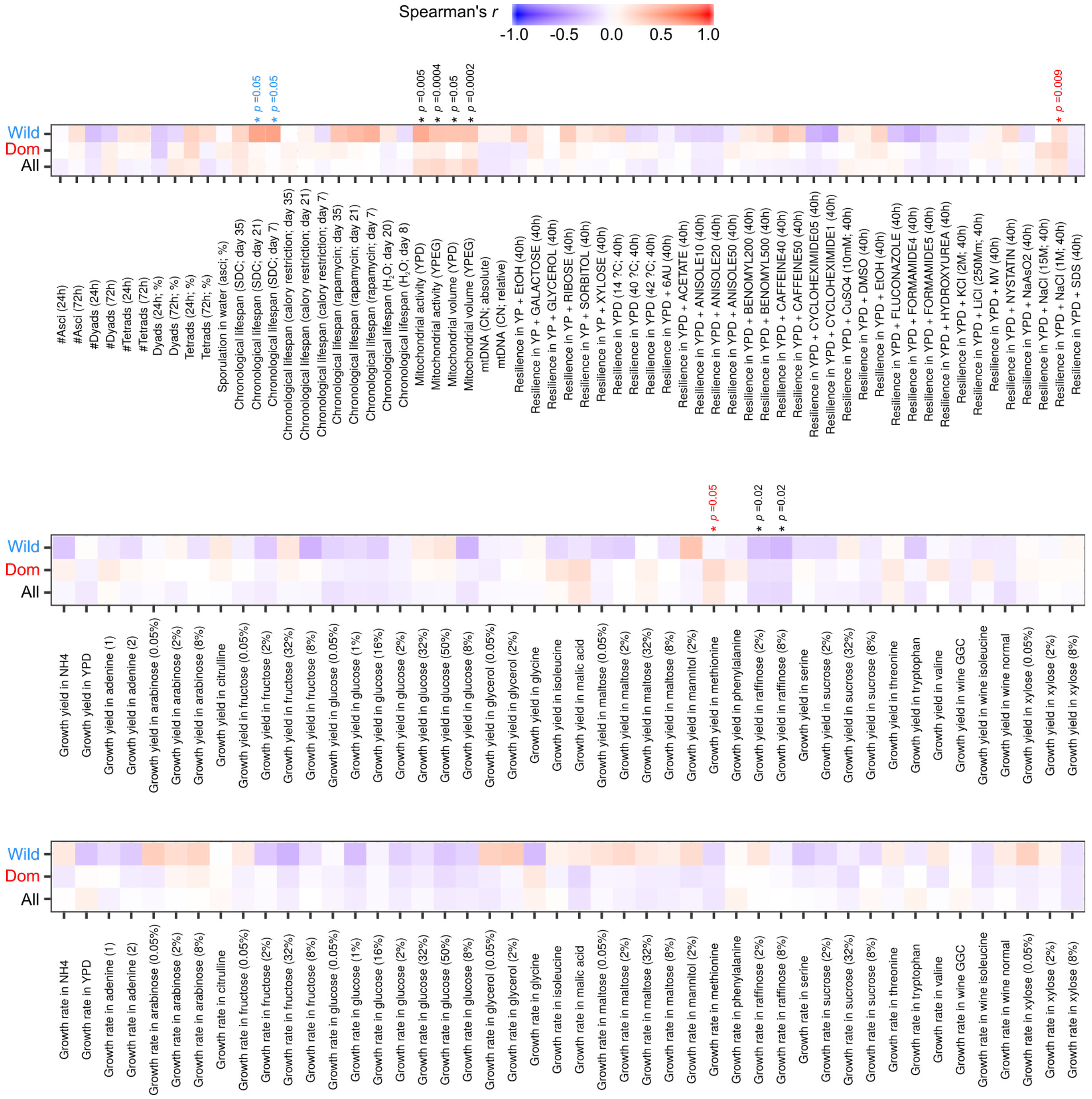


**Supplementary Figure 12 – TL and fitness traits.** Heatmap depicting the Spearman's correlation coefficient between telomere length and other phenotypes (*n*=155). Blue tones represent negative coefficients while red tones represent positive coefficients. Asterisks indicate phenotypes whose association with telomere length is statistically significant (two-tailed Wilcoxon test and FDR correction for multiple hypothesis testing), and their colour represents the group for which the association is significant (all, domesticated or wild isolates). *p*-values are indicated on top of each asterisk.
